# ASTRA: a deep learning algorithm for fast semantic segmentation of large-scale astrocytic networks

**DOI:** 10.1101/2023.05.03.539211

**Authors:** Jacopo Bonato, Sebastiano Curreli, Sara Romanzi, Stefano Panzeri, Tommaso Fellin

## Abstract

Changes in the intracellular calcium concentration are a fundamental fingerprint of astrocytes, the main type of glial cell. Astrocyte calcium signals can be measured with two-photon microscopy, occur in anatomically restricted subcellular regions, and are coordinated across astrocytic networks. However, current analytical tools to identify the astrocytic subcellular regions where calcium signals occur are time-consuming and extensively rely on user-defined parameters. These limitations limit reproducibility and prevent scalability to large datasets and fields-of-view. Here, we present Astrocytic calcium Spatio-Temporal Rapid Analysis (ASTRA), a novel software combining deep learning with image feature engineering for fast and fully automated semantic segmentation of two-photon calcium imaging recordings of astrocytes. We applied ASTRA to several two-photon microscopy datasets and found that ASTRA performed rapid detection and segmentation of astrocytic cell somata and processes with performance close to that of human experts, outperformed state-of-the-art algorithms for the analysis of astrocytic and neuronal calcium data, and generalized across indicators and acquisition parameters. We also applied ASTRA to the first report of two-photon mesoscopic imaging of hundreds of astrocytes in awake mice, documenting large-scale redundant and synergistic interactions in extended astrocytic networks. ASTRA is a powerful tool enabling closed-loop and large-scale reproducible investigation of astrocytic morphology and function.

## Introduction

Astrocytes tile the entire central nervous system in non-overlapping domains ^1^ interacting with neurons, vasculature, and other glial cells. Astrocytes exhibit a form of excitability based on changes in the intracellular calcium concentration ^2–4^. These calcium signals can be tightly related to synaptic activity ^5–7^ and to sensory inputs ^8–11^ and are instrumental for cognitive performance ^12, 13^. More recently, astrocytic calcium signals have been shown to encode information about external variables in awake behaving animals ^14–17^.

Astrocytic calcium signals can be monitored with high spatial resolution in the intact brain of awake animals using two-photon microscopy and chemical or genetically encoded calcium (GECI) indicators ^10, 18, 19^. The spatial features of astrocytic calcium signals are inextricably related to their highly ramified morphological structure, with thin (µm- and sub-µm-size) processes stemming from the soma and covering a tissue volume of ∼ 70-100 µm diameter (astrocytic domain). Within this tissue volume, astrocytic processes contact few neural cell bodies, hundreds of dendrites, and thousands of synapses (Halassa, Fellin, and Haydon 2007). Astrocytic calcium dynamics that can be localized to specific subcellular compartments including the cell body and portions of processes ^20–22^, can have different temporal characteristics ^10, 19, 20, 22–24^, and be coordinated across multiple astrocytes spanning hundreds, potentially thousands, of µm of brain tissue ^19, 23, 25, 26^.

Because of these complex properties, it is important to have software tools that systematically identify process and soma in two-photon functional recordings. Methods to identify and semantically segment astrocytic subcellular regions displaying calcium dynamics in individual astrocytes such as GECI-Quant ^19^ and CHIPS ^27^ are available. However, they heavily depend on data acquisition conditions, require the user to arbitrarily set several parameters, and need significant computation time. Other approaches identify calcium events within and across astrocytes as coherent, spatially-confined activity regions, based on pixel-wise fluorescence dynamics ^4, 28–31^. These event-based approaches are computationally demanding, still require *a posteriori* segmentation to relate identified events to astrocytic morphology, and have not been validated on large fields-of-view comprising large networks of astrocytes. As a result, currently available approaches still do not allow a fully automated, reproducible, fast and scalable analysis of astrocytic calcium signals within individual cells and across large populations that generalizes well to unseen datasets, different indicators, and experimental parameters. Therefore, developing fast, automated, and reliable image segmentation methods to analyze large-scale astrocytic calcium signals is of utmost urgency.

Similar challenges are faced in neuronal calcium imaging, where most advanced neural segmentation methods include both unsupervised and supervised machine learning approaches ^32–38^. However, approaches specifically developed for segmentation of neuronal calcium imaging t-series cannot be readily applied to the analysis of astrocytic calcium signals, as demonstrated in this work, because the spatial and temporal features of astrocytic calcium signals are intrinsically different from those of neurons.

Here we present ASTRA, Astrocytic calcium Spatio-Temporal Rapid Analysis, a novel deep learning software that performs fast, precise, scalable, and fully automated semantic segmentation of astrocytic two-photon imaging t-series. ASTRA combines feature engineering and a deep learning algorithm to enable scalable and repeatable analysis. We validated ASTRA using the consensus annotation generated by three human experts of four novel two-photon microscopy datasets recorded in awake head-fixed animals. These annotated datasets are shared here, for future community-based development and benchmarking of algorithms for the detection and segmentation of astrocytes. ASTRA performed cell detection (identification of somata of astrocytes) and semantic segmentation (identification and labeling of cell soma and proximal processes) with near-human-expert performance. ASTRA outperformed all tested state-of-the-art software for the analysis of astrocytic and neuronal signals, was endowed with features combining segmentation with event-based analyses to identify astrocytic cellular domains, and generalized well across indicators and acquisition conditions. ASTRA also scaled well to large datasets, allowing rapid automated analyses of entire databases characterized by many recording sessions and enabled analysis of the first report of simultaneous functional imaging of hundreds of astrocytes distributed over millimeters of cortical tissue recorded in awake mice using two-photon fluorescence mesoscopic imaging.

## Results

### ASTRA: structure and analysis workflow

Here we developed ASTRA, a software that combines statistical image analysis and deep learning to perform fully automated segmentation of astrocytes imaged with two-photon fluorescence microscopy. ASTRA operates on astrocytic two-photon imaging t-series and uses both morphological and dynamical information to provide, as output, three classes of regions of interest (ROIs): somata, processes, and cross-correlated regions denoting a two-dimensional measurement of the astrocytic domains (Fig, 1).

ASTRA includes a training pipeline and an inference pipeline (Fig. 1A-B, Fig. S1A-B). Each pipeline analyzes a dedicated training and inference dataset, respectively. The training set is the subset of the data (e.g., a subset of the available fields-of-view (FOVs)) annotated by human experts. The training set is used to optimize ASTRA’s pre-processing hyper-parameters and train the weights of the Deep Neural Network (DNN), which performs semantic segmentation. The inference dataset is a separate data subset (e.g., FOVs not included in the training set), on which the algorithm performs inference (i.e., semantic segmentation). Because pre-processing parameters and DNN weights are optimized automatically on the training dataset, the inference pipeline runs on test data without human supervision.

**Figure 1.**
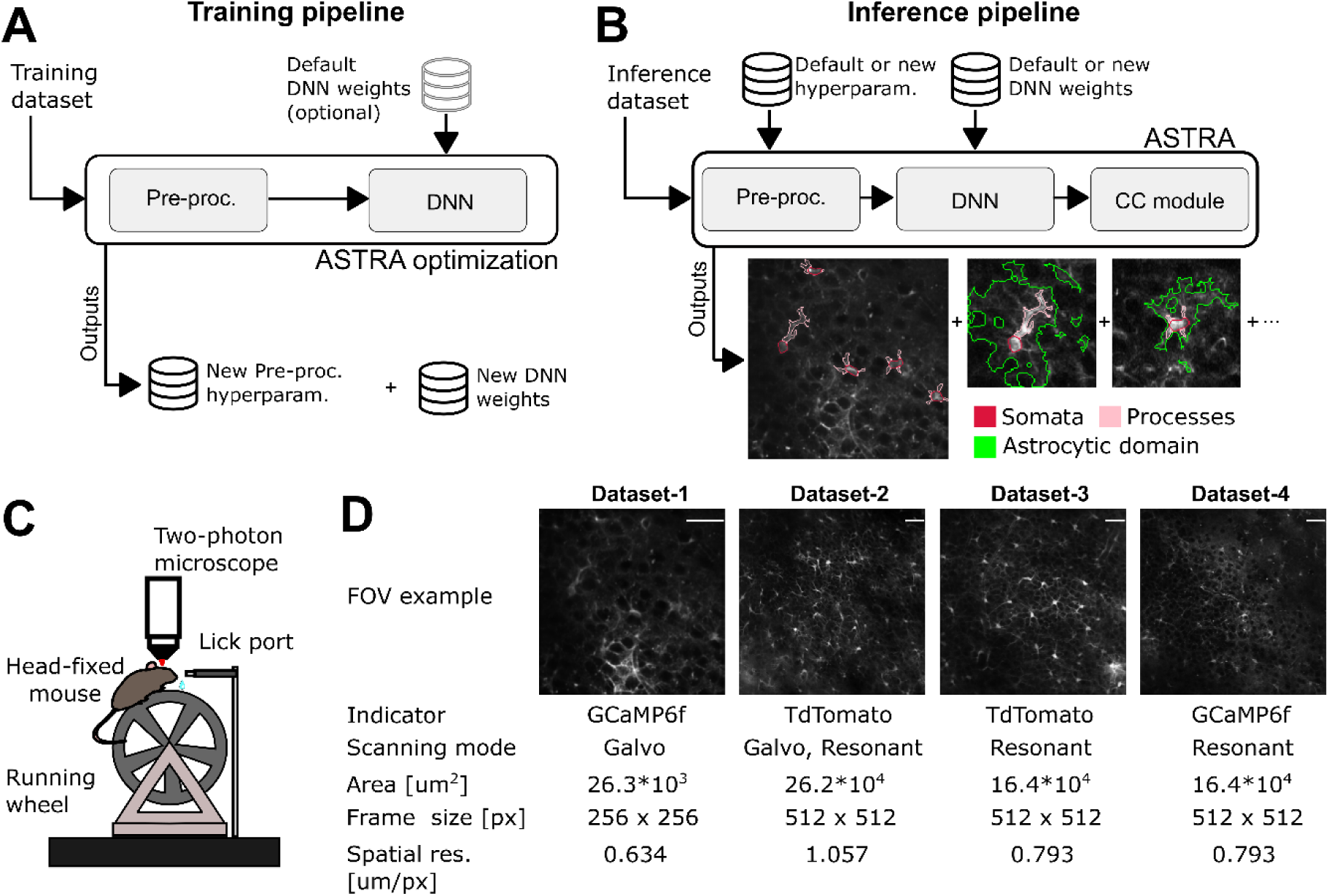
ASTRA: a machine learning algorithm for fast and automated semantic segmentation of astrocytes. A-B) Flow-charts of ASTRA segmentation pipelines for training (A) and inference (B). At the end of the training pipeline, pre-processing hyperparameters and DNN weights are saved. At the end of the inference pipeline spatial coordinates corresponding to somata, processes, and cross-correlated regions are saved. C) Two-photon Ca^2+^ imaging of hippocampal astrocytes was performed in head-fixed mice running on a wheel. D) Four datasets were initially used for ASTRA training and testing. Details of each dataset are listed in the figure. Each dataset was manually segmented by 3 expert annotators. White bar on the top-right of each image represent the scale bar. Dataset-1, 40 µm, Dataset-2, 50 µm Dataset-3, 40 µm, and Dataset-4, 40 µm.

The inference pipeline comprises three main steps: i) *pre-processing*; ii) *semantic segmentation*;iii) *subcellular cross-correlation analysis* (Fig. 1B and S1B). The *pre-processing* step computes a bi-dimensional reconstruction of the recorded field-of-view (FOV), compressing spatial and temporal features into a highly informative spatiotemporal projection (Fig. S2A-B, see also Methods). Pre-processing enhances astrocytic subcellular structures (e.g., processes and somata) and decreases acquisition noise from the bi-dimensional reconstruction of the FOV. The spatial component of the pre-processing uses histogram equalization and kernel convolution to compute a sharpened spatial map (Fig S2A, right) from the median projection of the time series, enhancing astrocytic morphological substructures. Then, it evaluates statistically pixel-wise temporal dynamics to set an optimal intensity threshold used to classify pixels that display foreground fluorescence. Foreground pixels are maintained in the sharpened spatial map while non-foreground (background) pixels are set to 0. The output of pre-processing feeds into the *Segmentation* step, which performs subcellular semantic segmentation of astrocytic somata and processes using U-Net-^39^ based DNN architecture (Fig. S1E). The training of the DNN weights on the annotated training datasets becomes feasible on the relatively limited data size typical of conventional two-photon imaging datasets because of the efficient feature-engineering during the pre-processing, and because ASTRA takes advantage of transfer-learning by employing a DNN encoder ^38, 40–42^ pre-trained on ImagNet dataset ^43^ (see Methods). Finally, the *subcellular cross-correlation analysis* identifies regions of the astrocytic domain showing fluorescence signals that are statistically correlated to the ones present in the semantically segmented regions of the astrocyte (see Methods).

We provide ASTRA with default pre-processing hyperparameters and DNN weights trained extensively on two different two-photon microscopic datasets. With these parameters, the inference pipeline can readily operate even on previously unseen data, as extensively demonstrated below on several datasets. However, ASTRA also offers users a further-training pipeline (Fig. 1A), which allows the inclusion of new own training data annotated with ImageJ ^44^. This additional pipeline can be used to further refine (and export for future use) the DNN weights and the pre-processing parameters to optimize the software to the applications at hand.

The ASTRA inference pipeline works fast on either CPUs or GPUs. Retraining the DNN with new annotated data provided by the user can be done on a single GPU or in parallel on multiple GPUs, setting simple options in the code.

### Datasets for consensus annotation and algorithm training and benchmarking

To train and benchmark ASTRA, we recorded and analyzed four datasets of two-photon fluorescence hippocampal recordings in awake head-fixed mice running on a wheel (Fig. 1C). The four datasets (Fig. 1D) differed for the type of fluorophore which was expressed in astrocytes (e.g., GCaMP6f and Td-Tomato), imaged area (from 26.3 x 10^3^ µm^2^ to 26.2 x 10^4^ µm^2^), acquisition procedures (galvanometric mirror-based imaging *vs* resonant scanning imaging), and pixel resolution (from 0.63 µm/ pixel to 1.06 µm/ pixel).

We generated a manual consensus annotation of all four datasets. Three expert annotators detected and segmented astrocytes, labelling somata and individual processes. Annotators had access to both the raw t-series and the bi-dimensional projections of the t-series obtained using the spatial component of the pre-processing pipeline. Annotators detected astrocytes on the t-series, while segmenting subcellular structures on bi-dimensional projections. After each expert annotator labeled independently the data, annotators were asked to converge on a consensus by resolving each single annotation discrepancy according to standard procedures (Methods). The result of this procedure was termed “consensus annotation” (Fig. 2A, Fig. S3). Consensus annotation was used, following standard practice ^36, 37^, to train the algorithm and to benchmark its performance (Fig. 2B). The four datasets, including individual and consensus annotations, will be shared upon publication.

**Figure 2.**
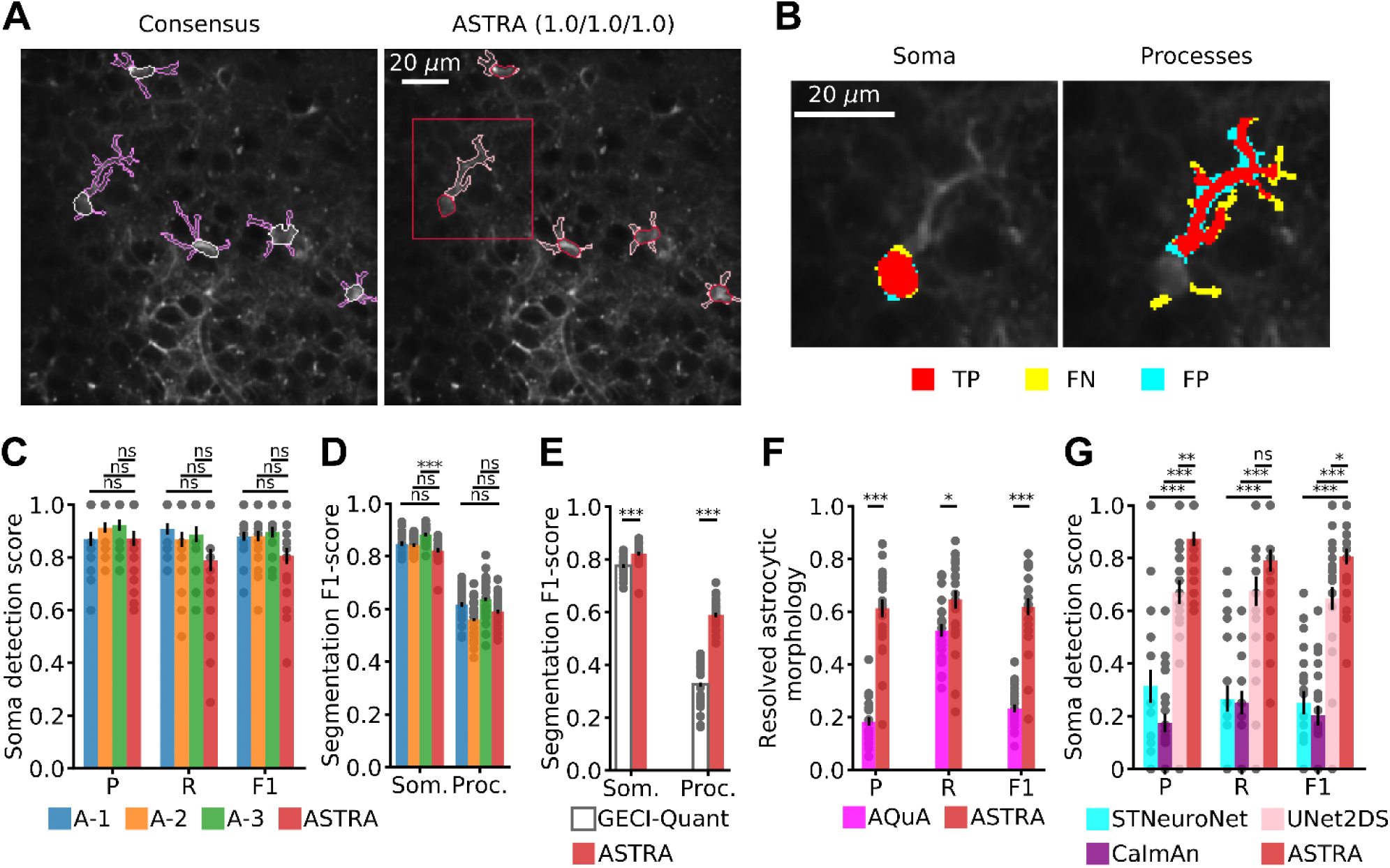
Evaluation and benchmarking of ASTRA on dataset-1. A) Representative comparison of consensus annotations (left, FOV id: 2) and ASTRA semantic segmentation (right). On the top of the right image are reported somata detection Precision, Recall, and F1-score for FOV id: 2. B) Representative example of the comparison of somata and processes segmentations between ASTRA and the consensus annotations. True positive pixels (red), false negative pixels (green), and false positive (cyan) are shown. C) Performance of the three annotators A-1, A-2, and A-3 against ASTRA. Precision (P), Recall (R), and F1-score (F1) are shown. Two-sided Wilcoxon rank sum test N= 24; leave-one-out cross validation (LOOCV) results. In this as well as in following figures: n.s., not significant, *P < 0.05, **P < 0.005 and ***P < 0.0005. D) F1-score for somata and processes segmentation for annotators and ASTRA. Two-sided Wilcoxon rank sum test, N= 24, LOOCV results. E) F1-score for somata and processes segmentation of GECI-Quant and ASTRA. Two-sided Wilcoxon rank sum test N= 24; LOOCV results. F) Astrocytic morphology reconstructed using ASTRA segmentations and AQuA event detection. Two-sided Wilcoxon rank sum test N= 24; LOOCV results. G) Soma detection performance of STNeuronet, CaImAn, UNet2DS, and ASTRA. Two-sided Wilcoxon rank sum test N= 24; LOOCV results. See also table S1, S2, and S3.

We first utilized the consensus annotation to quantify the performance and consistency of human experts (Fig. 2C-D). Somata detection performance of human annotators was highly accurate (high F1-score, Fig. 2C), demonstrating a high human consistency on astrocyte somata detection. Conversely, segmentation performance of human annotators showed lower F1-scores (Fig. 2D). This was especially true for processes (Fig. 2D), implying that annotation by a single human grader can be unreliable (Supplementary information Tab. S1, Tab. S4, Tab. S5, and Tab. S6), and that benchmarking and training should be better done using the consensus annotation ^36, 37^.

### ASTRA achieves human performance and replicates previously published results obtained with manual annotation

We then used the consensus annotation datasets to train and test ASTRA. We first used dataset-1, which comprises a set of 24 two-photon calcium imaging recordings of the CA1 hippocampal region recorded in head-tethered awake mice, which were spontaneously walking on a wheel (Fig. 1C). In the hippocampal CA1 region, astrocytes expressed the genetically encoded calcium indicator GCaMP6f after adeno-associate viral transduction. GCaMP6f signals were collected from a FOV of area approximately 26 x 10^3^ µm^2^ with a spatial sampling of 0.634 µm/pixel (Fig. 1D, see Methods).

We tested ASTRA on dataset-1 using leave-one-out cross-validation (see Methods). Fig. 2A and 2B show an example of annotations obtained by ASTRA on a representative FOV compared to the consensus annotation. Importantly, Precision, Recall, and F1-score of ASTRA detection were high and did not significantly differ from those of the annotators (Fig. 2C, SI Tab. S1). Segmentation F1-score was high for somata and similar to that of two out of the three annotators (Fig. 2D, S4A-B, SI Tab. S2). Segmentation F1-score was lower for processes, but similar to that of all annotators (Fig. 2D, Fig. S4A-B, SI Tab. S2). Overall, these results indicate that ASTRA detection and segmentation accuracy levels are comparable to those of individual human experts.

Given that ASTRA performed like human experts, we tested if it could replicate, in a fast and fully automated way, previously published results based on manual segmentation of astrocytes. We applied ASTRA to perform automated semantic segmentation of CA1 hippocampal astrocytes imaged with two-photon functional microscopy during mouse virtual navigation (^15^, Fig. S5). Astrocytic ROIs detected using ASTRA resembled human detections used in ^15^ (mean ± sem, Precision: 0.86 ± 0.05, Recall: 0.75 ± 0.07, F1: 0.78 ± 0.05, N = 7 imaging sessions from 3 mice). Using ASTRA annotations, we were able to replicate the major results described in (^15^, Fig. S5 C-E), demonstrating that astrocytic spatial tuning parameters obtained by manual annotation were recapitulated using ASTRA semantic segmentation (Fig. S5 F-G). Importantly, while manual annotation of the dataset described in ^15^ required several days of work, ASTRA segmented the whole dataset in ^15^ in approximately 9 minutes without human intervention or arbitrary parameter settings. ASTRA can thus be used for fast, automated, and reproducible analysis of entire datasets and compares well with manual expert annotation of the same datasets.

### ASTRA outperforms state-of-the-art algorithms for the analysis of astrocytic and neuronal signals

We benchmarked ASTRA against several major methods for analysis of two-photon fluorescence recordings of astrocytes and neurons (Fig. 2E-G, Fig. S4E-L).

We first compared ASTRA with GECI-Quant ^19^, a threshold-based user-supervised software for the analysis of astrocytic two-photon calcium imaging data. For each FOVs, one of our annotators (annotator A-1) manually identified astrocytic domains and defined the intensity thresholds to segment somata and processes (see Fig. S4F). This manual input is needed to run GECI-Quant. The indications of the GECI-Quant documentation ^19^ were closely followed during this procedure. The detection F1-score of GECI-Quant was not significantly different from that of ASTRA (two-sided Wilcoxon rank sum test N=24, Fig. S4G), not surprisingly because the domain identification of GECI-Quant was performed by a human expert and ASTRA performed as a human expert. However, segmentation performances of GECI-Quant were lower than those of ASTRA for somata and especially so for processes (Fig. 2E and Fig. S4H-I, SI Tab. S2, two-sided Wilcoxon rank sum test, N= 24).

We then compared the performance of ASTRA to that of AQuA ^28^, an event-based algorithm which identifies ROIs associated with astrocytic calcium events based on the coherence of fluorescence dynamics across pixels. Although the AQuA definition of events does not consider morphological constraints, we reasoned that it should be possible to use AQuA to potentially identify astrocytic somata and processes, because a subsets of calcium events should be restricted to astrocytic soma or processes. We thus identified the morphology of putative somas and processes by thresholding a time-averaged spatial map of calcium events identified by AQuA, and we compared it to the consensus annotation. The segmentation so obtained with AQuA had precision, recall, and F1-score against consensus annotation lower than ASTRA’s (Fig. 2F, SI Tab. S3, two-sided Wilcoxon rank sum test, N= 24). Taken together, these results demonstrate that ASTRA outperforms the tested state-of-the-art methods used for the analysis of astrocytes data in identifying astrocytic somata and processes.

We then asked whether software for segmentation of neurons from two-photon imaging can be used for the segmentation of astrocytes. We compared ASTRA with STNeuroNet ^37^, UNet2DS ^39^, and CaImAn ^36^, three state-of-the-art algorithms, which perform binary classification (foreground *vs* background) of pixels in FOVs to identify neuronal ROIs. STNeuroNet and UNet2DS use DNN in a way conceptually comparable to ASTRA, but are specialized for neurons. In contrast, CaImAn is a fully unsupervised algorithm not based on deep learning. When initially applying to astrocytic data STNeuroNet and UNet2DS in their off-the-shelf form, they almost never identified regions labeled as astrocytic soma or processes in the consensus (data not shown). We thus retrained the weights of STNeuroNet and UNet2DS on our astrocytic consensus data. Moreover, we adjusted the pre- and post-processing steps of STNeuroNet to constraint source detection using parameters based on astrocytic (rather than neural) calcium dynamics and morphology (Methods). We instead straightforwardly applied CaImAn without making any change, as it is a fully unsupervised algorithm. We found that all three neural algorithms identified only regions that were labeled as soma in the consensus, but they did not detect regions labeled as processes in the consensus (Fig. S4J-L). This was not surprising because these neural algorithms were conceived to mostly detect neuronal cell somata. We thus analyzed the output of these algorithms only considering astrocyte somata detection (Fig. 2G, SI Tab. S2). We found that the F1-score of somata detection performance of UNet2DS (mean ± sem, 0.65 ± 0.04, N = 24 was significantly higher than that of CaImAn (mean ± sem, 0.20 ± 0.04, N = 24) and STNeuroNet (mean ± sem 0.27 ± 0.05, N = 24, two-sided Wilcoxon rank sum test). Importantly, somata detection performance of all three neural algorithms was inferior to that of ASTRA (Fig. 2G, two-sided Wilcoxon rank sum test, N = 24).

Together, these results stress the need to introduce dedicated algorithms for astrocytic segmentation and indicate that ASTRA outperforms available analysis methods specifically developed for neuronal datasets, even after adjusting them to astrocytic analysis.

### Identification of functional domains of individual astrocytes using ASTRA

Thin (diameter < 1 µm) astrocytic processes substantially contribute to fill the domain of brain tissue occupied by a single astrocyte (astrocytic domain) and display information-rich calcium dynamics ^19–22, 30^. However, the identification of these thin structures is challenging, because of the dimension of thin astrocytic processes is smaller than the spatial resolution of two-photon microscopy. The difficulty in identifying thin astrocytic processes makes it challenging to measure the astrocytic domain based only on morphological features. We thus implemented within ASTRA an algorithm based on activity correlation measurement, termed “subcellular cross-correlation analysis”, to reproducibly identify, based on activity measurements, the extent of an astrocytic “functional” domain, including somata, main processes, and subresolved cellular compartments. This analysis automatically selected pixels within the typical extent of a domain of an individual astrocyte. Based on previous reports ^1, 45^, we set the astrocytic domain as a circular region of radius ∼40 μm centered on the center of mass of the astrocyte soma. The fluorescence dynamics of the domain pixels were correlated to the pixels belonging to the semantically segmented ROIs (either somata or processes) of that same astrocyte (Fig. 3A). The output of this analytical procedure was a ROI of correlated pixels (Fig. 3A), which included cell somata and processes and which resembled anatomically defined astrocytic domains ^1, 45^.

**Figure 3.**
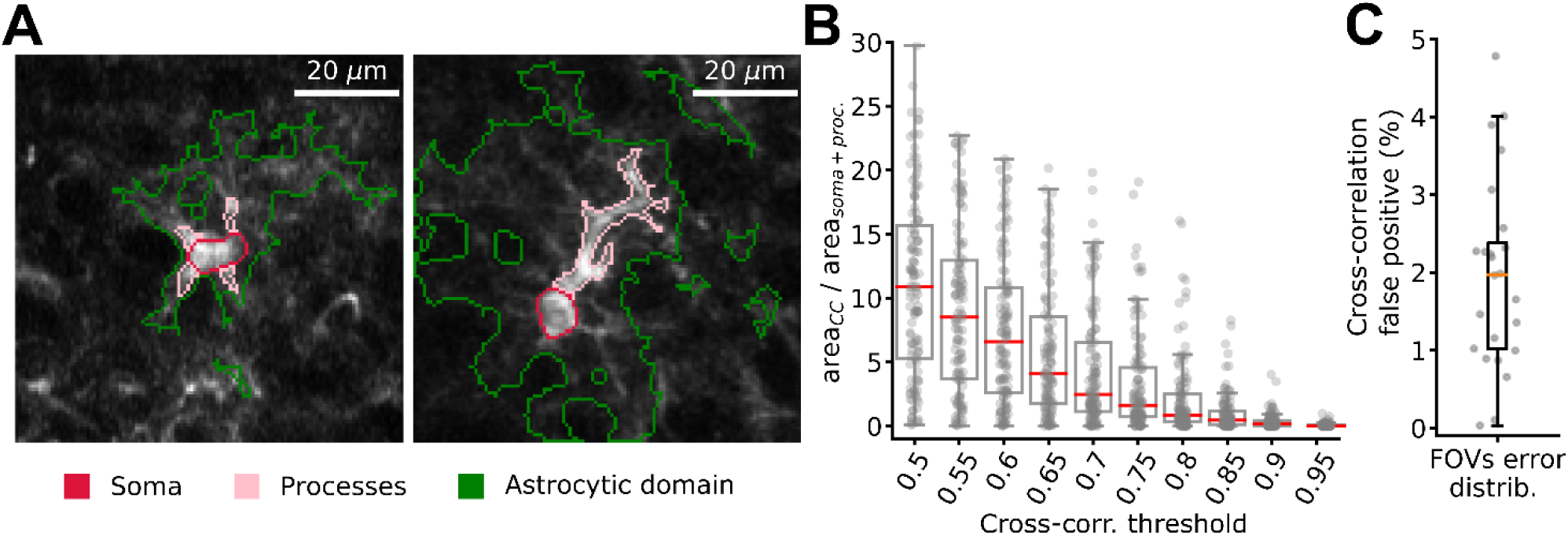
Identification of correlated calcium signals in astrocytic domains using ASTRA. A) Two representative examples of statistically correlated regions in the astrocytic domain identified with the cross-correlation module (FOV (Id:2)). ROIs corresponding to somata and processes are colored in red and pink, respectively. ROIs extracted using cross correlation are shown in green. B) Ratio of ROI area extracted using the cross-correlation module and ROI area obtained by summing soma and processes ROIs together as a function of the cross-correlation threshold. C) Cross correlation error distribution. The cross-correlation error was estimated as the mean percentage of false-positive pixels selected in each FOV sampling 1000 pixels outside astrocytes domains, which were not used to tune the cross-correlation threshold. Two-sided Wilcoxon rank sum test, N=24.

The identified astrocytic domain depended on a single parameter, the value of the cross-correlation threshold (Fig. 3B). Low threshold values selected larger areas, including potentially pixels belonging to neuronal structures (i.e., neuronal cell somata and processes). Conversely, high thresholds select smaller areas, possibly neglecting meaningful astrocytic structures. To set an optimal, intermediate, threshold value, we programmed ASTRA to dynamically auto-tune the cross-correlation threshold separately for each FOV, to control for the ratio of false positives. Once segmentation of the entire dataset was completed by ASTRA, the cross-correlation module randomly sampled 250 pixels located outside the astrocytic domains identified around the segmented astrocytic somas. ASTRA computed the cross-correlation between the activity of the randomly sampled pixels and the pixels inside ASTRA-segmented ROIs and estimated the false positive rate as the fraction of randomly sampled pixels with correlation above the set threshold. The algorithm then tested a grid of threshold values and automatically set the threshold as the smallest threshold value with false positive percentage error < 5% (see Methods). ASTRA then randomly sampled 1000 pixels located outside the astrocytic domains identified around the segmented astrocytic somas and used them to confirm that the false positive rate < 5 %. This procedure was effective at minimizing false positives. On dataset-1 and across all FOVs, this procedure selected pixels with an average false positive percentage of 2.0 ± 0.2 % (mean± sem, Fig. 3C). On dataset-1 the cross correlated area was 585 ± 57 µm^2^ (mean ± sem).

Importantly, the functional domains of individual astrocytes identified by ASTRA can then be used to seed the event-based analysis performed by AQuA ^28^. This was demonstrated in (Fig. S6), where we show examples of astrocytic domain identified by ASTRA, which were used as priors to instruct cell-specific AQuA analysis.

Taken together, these findings demonstrate that ASTRA could be used to identify functional domains of individual astrocytes encompassing the cell somata, main processes, and thin astrocytic structures. Moreover, combining ASTRA with the event-based analysis software AQuA allowed overlaying anatomical with functional analysis of astrocytic domains, enabling the extraction of previously hidden morpho-functional information from individual astrocytes recorded in two-photon GCaMP imaging experiments.

### ASTRA performance across signal-to-noise ratios

To investigate the performance of ASTRA as a function of the signal-to-noise ratio of two-photon images, we performed a set of comparative analyses on t-series from dataset-1 in which we artificially increased and decreased the peak signal-to-noise ratio (PSNR) of the fluorescent signals (see Fig. 4A). Manipulations ranged from nearly halving to nearly doubling the PSNR, with respect to the original data. ASTRA detection F1-score significantly decreased when the PSNR was strongly reduced (Fig. 4B, two-sided Wilcoxon rank sum test N = 24). However, the F1-score for the segmentation of somata and processes remained unaltered (Fig. 4C, Fig. S7A-B). These results showed that ASTRA semantic segmentations was robust to the degradation of the PSNR. In contrast, an increase of the PSNR resulted in an improvement of the F1-score for detection (Fig. 4B, two-sided Wilcoxon rank sum test N = 24) and of the F1-score for segmentation of processes (Fig. 4C, Fig. S7A-B; two-sided Wilcoxon rank sum test N = 24), with no significant change of the F1-score for somata segmentation.

**Figure 4.**
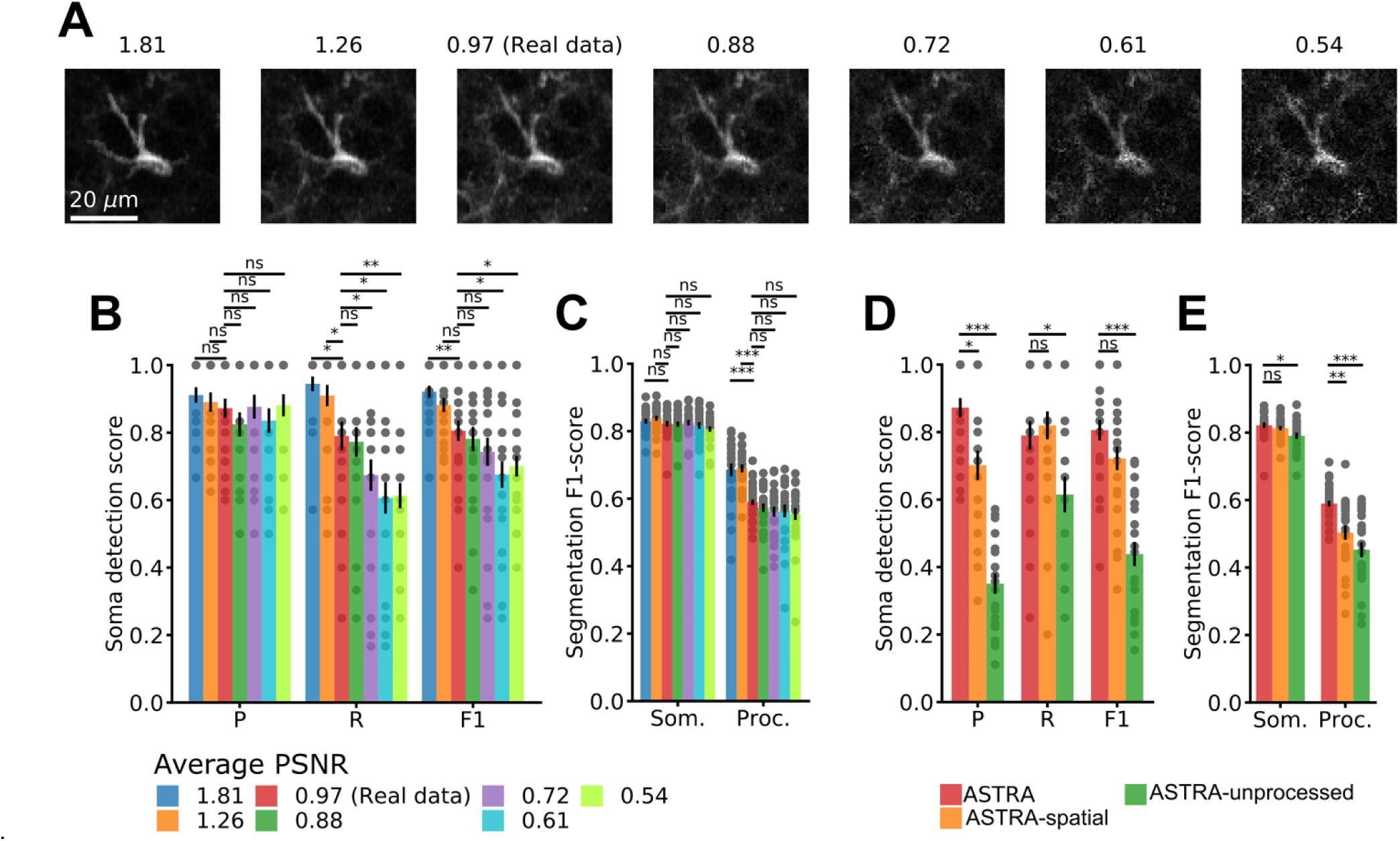
Impact of image noise and pre-processing on ASTRA performance. A) Representative image (single cell in FOV id 2) under various simulated noise regimes. Values of peak signal-to-noise ratio (PSNR) for each noise regime are reported above the images. B) Precision, Recall, and F1-score for soma detection performance for different PSNRs. Two-sided Wilcoxon rank sum test, N = 24; LOOCV results. C) F1-score for segmentation of somata and processes across different PSNRs. Two-sided Wilcoxon rank sum test, N = 24; LOOCV results. D-E) ASTRA detection and segmentation performance as a function of the omission of ASTRA pre-processing steps. We omitted either the temporal pre-processing step (ASTRA-Spatial) or all the pre-processing steps (ASTRA-unprocessed). Soma detection Precision, Recall, and F1 are reported in D. Two-sided Wilcoxon rank sum test, N = 24; LOOCV results. The segmentation F1-score for somata and processes are shown in E. Two-sided Wilcoxon rank sum test, N = 24; LOOCV results.

We also evaluated the other state-of-the-art detection and segmentation methods, described above, on these artificial datasets with modified PSNR. We first tested GECI-Quant detection and segmentation performance under high PSNR conditions (1.81 times the original PSNR, Fig. S7C-G). For each FOVs one annotator manually defined astrocyte somatic regions, astrocyte domains, and intensity thresholds (Fig. S7C). We found that the detection F1-score of GECI-Quant was significantly lower than that of ASTRA (Fig. S7D, two-sided Wilcoxon rank sum test N = 24). GECI-Quant F1-scores for process segmentation was also lower than that of ASTRA (Fig. 7G, two-sided Wilcoxon rank sum test N = 24). We then tested algorithm developed for the analysis of neuronal datasets. We found that STNeuroNet and CaImAn showed lower performance across all PSNR conditions when compared with ASTRA (Fig. S7 H-I, two-sided Wilcoxon rank sum test on all artificial datasets, N = 24, compared with real data). UNet2DS showed lower F1-score compared to ASTRA, but this decrease was significant only for PSNR = 0.88 (Fig. S7J, two-sided Wilcoxon rank sum test N = 24). The stability of UNet2DS to changes in PSNR can be understood considering that UNet2DS use only the mean projection in time of the recorded videos and that the injected Gaussian noise does not affect this projection.

Overall, these results show that ASTRA performance remains stable even with low PSNR, and favorably compares with the performance of state-of-the-art methods for the analysis of astrocytic and neuronal functional signals over a wide range of PSNR.

### Pre-processing is important for ASTRA performance

The ASTRA DNN segmentation operated on fluorescence imaging t-series downstream of the pre-processing modules, which performed image processing to enhance spatial information and performed temporal processing to filter out background from foreground signals. We produced two reduced versions of ASTRA. A first version called (ASTRA-unprocessed, Fig. 4D-E and Fig. S8) in which we removed both spatial and temporal pre-processing by performing DNN analysis directly on the raw median projection of the t-series. A second reduced version (ASTRA-spatial, Fig. 4D-E and S8) where we only removed the temporal pre-processing. Compared against the consensus annotation, we found that ASTRA-unprocessed had considerably lower performance than the full version of ASTRA (Fig. 4D-E, two-sided Wilcoxon rank sum test N = 24) for somata detection and for somata and process segmentation. Moreover, we observed that ASTRA-spatial had similar performance for somata detection and segmentation, but much lower performance for process segmentation than the full version of ASTRA (Fig. 4D-E, two-sided Wilcoxon rank sum test N = 24). These findings highlight the importance of the pre-processing step for ASTRA performance.

### Number of frames needed to reach good performance when training ASTRA from scratch

Although we trained and tested ASTRA using all experimentally recorded imaging frames in each dataset, we wanted to estimate how ASTRA would have performed had we had less recorded frames. We thus repeated our analyses after decimating dataset 1 to only include between 50 and 400 frames, rather than the 550-750 frames of the original t-series. This is of interest because the size of t-series can greatly vary across experiments in two-photon imaging experiments. The ASTRA detection F1-score remained stable as long as the t-series was longer than 200 frames (Fig. S9A, two-sided Wilcoxon rank sum test N = 24) and the F1-score for process segmentation also remained unchanged for t-series longer or equal than 100 frames (Fig. S9D, two-sided Wilcoxon rank sum test, N = 24). These results suggest that 100-200 frames per FOV are sufficient to train ASTRA.

### ASTRA generalizes across indicators and acquisition parameters

To investigate whether ASTRA generalizes across experimental preparations, acquisition parameters, as well as to never-seen-before data, we benchmarked it on datasets 2-4.

We first investigated whether ASTRA could be trained anew on a novel dataset with very different characteristics. We thus trained anew and tested ASTRA (using exactly the same procedure detailed above for dataset-1) on the dataset-2, a set of eight two-photon imaging recordings collected in either resonant- or galvanometric mirror-based scanning in the hippocampus of head-fixed awake animals spontaneously walking on a wheel (Fig. 1C-D). In this dataset, hippocampal astrocytes specifically expressed TdTomato and fluorescence signals were collected from a FOV of area 26.2 x 10^4^ µm^2^ with pixel size of 1.06 µm/pixel (Fig. 1D, see Methods). On dataset-2, ASTRA detection and segmentation performance reached the level of the individual human experts (Fig. S10C-F, Tab. S4). This result suggests that ASTRA can be readily trained to reach human expert performance, regardless of the indicator and of the acquisition parameters used.

We then tested whether ASTRA can be used with the pre-trained DNN weights and without any further training on never-seen-before datasets with different indicators and acquisition parameters. We thus took ASTRA with the DNN pre-trained weights (obtained by training on dataset 1 and available as default weights in the online ASTRA software) and we applied it straightforwardly to two new never-seen-before datasets (dataset-3 and dataset-4). Dataset-3 was composed of a set of seven two-photon imaging recordings collected in resonant scanning mode in the hippocampus of head-tethered awake animals spontaneously walking on a wheel (Fig. 1 C-D). Hippocampal astrocytes specifically expressed TdTomato and fluorescence signals were collected from a FOV of area 16.4 x 10^3^ µm^2^ with a pixel size of 0.79 µm/pixel (Fig. 1D). Dataset-4 included a set of ten two-photon calcium imaging t-series collected in the resonant scanning modality in head-fixed awake animals free to run on a wheel (Fig. 1C-D). In dataset-4, hippocampal astrocytes specifically expressed GCaMP6f and fluorescence signals were collected from a FOV of area 16.4 x 10^3^ µm^2^ with a pixel size of 0.79 µm/pixel (Fig. 1D). Both dataset-3 and dataset-4 had a consensus annotation obtained as for dataset-1. Results of benchmarking against the consensus revealed that ASTRA reached the level of human experts for both dataset-3 (Fig S10G-L, SI Tab. S5) and dataset-4 (Fig. 5 and S11, see also SI Tab. S6). Thus, in the case of the two never-seen-before dataset-3 and dataset-4, ASTRA reached human expert performance with pre-trained weights, implying that there would be no benefit in re-training the DNN adding new consensus annotated data from the new datasets (something that we explicitly verified, data not shown).

**Figure 5.**
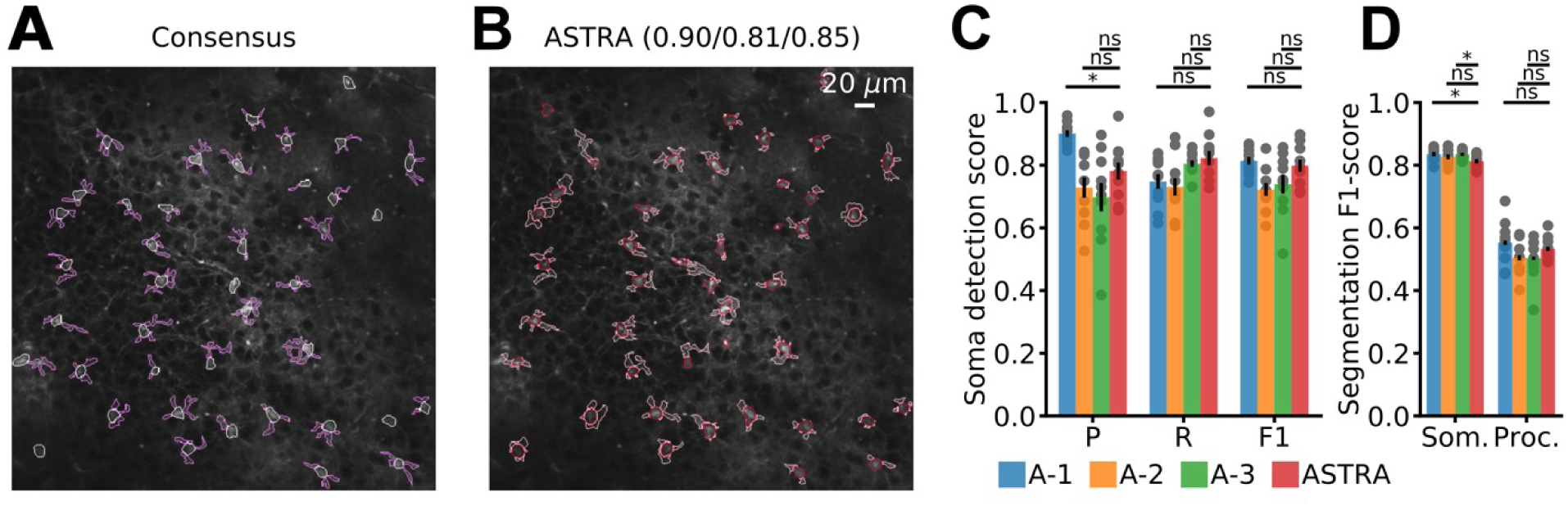
ASTRA performance on never-seen-before data. A) Consensus annotation of FOV (Id: 1, dataset-4). B) ASTRA segmentation on the same FOV shown in A. C) Soma detection performance is reported as Precision (P), Recall (R), and F1-score (F1) for the three human annotators (A-1, A-2, and A-3) and for ASTRA. Two-sided Wilcoxon rank sum test, N=10; LOOCV results. D) F1-score for segmentation of somata and processes for the three human annotators (blue, yellow, and green) and ASTRA (red). Two-sided Wilcoxon rank sum test, N=10; LOOCV results. See also table S5.

Overall, these results demonstrate that ASTRA compared to individual human annotators even on never-seen-before data.

Given the above success, however, it is conceivable that in some other never-seen-before datasets, ASTRA may not reach human expert performance when using off-the-shelf pretrained weights. In such case, ASTRA offers the possibility to fine tune the DNN weights by retaining on new data added by the users starting from the pre-trained weight that we made available with the software or from any other set of initial weights chosen by the user (Fig. 1A, Fig S12).

### Automated analysis of two-photon mesoscopic imaging of astrocytes using ASTRA

The activity of multiple astrocytes is known to be correlated over spatial scales of few hundreds of µm, which are typically imaged with two-photon microscopes (reviewed in ^4, 46, 47^). However, very little is known about how the activity of astrocytes is organized at the network level over regions spanning several millimeters. It is now possible to perform high-resolution functional imaging over distances of millimeters with two-photon large FOV microscopes (mesoscopes) ^48^. However, analysis of mesoscopic images requires the segmentation of hundreds of ROIs in each FOV, making manual annotation prohibitive. Thus, an important application of ASTRA is enabling analyses of mesoscopic FOVs with distributed astrocytic networks encompassing hundreds of cells. Here, we demonstrate the usefulness of ASTRA for this application.

To this aim, we performed for the first time two-photon mesoscopic imaging in awake head-fixed mice expressing GCaMP6f in cortical astrocytes (Fig. 6). Mice were free to run on a wheel and licked at will from a water spout. We ran ASTRA on these mesoscopic t-series. An example of mesoscopic FOV segmentation is shown in Fig. 6A. On average, ASTRA extracted, 119 ± 29 astrocyte somata *per* FOV, N = 15 FOVs (average area of FOV, ∼ 2.3 mm^2^). Moreover, we found that ASTRA identified processes that were less numerous (per identified soma), smaller, and shorter than those identified with standard two-photon microscopes analyzed above (see Fig. S13). This was most likely due to the lower numerical aperture of the two-photon mesoscope objective ^48^ with respect to that of standard two-photon microscope objectives, which implies lower spatial resolution in mesoscopic recording compared to standard two-photon recordings. We thus focused our next mesoscopic analyses on networks of astrocytic somata.

**Figure 6.**
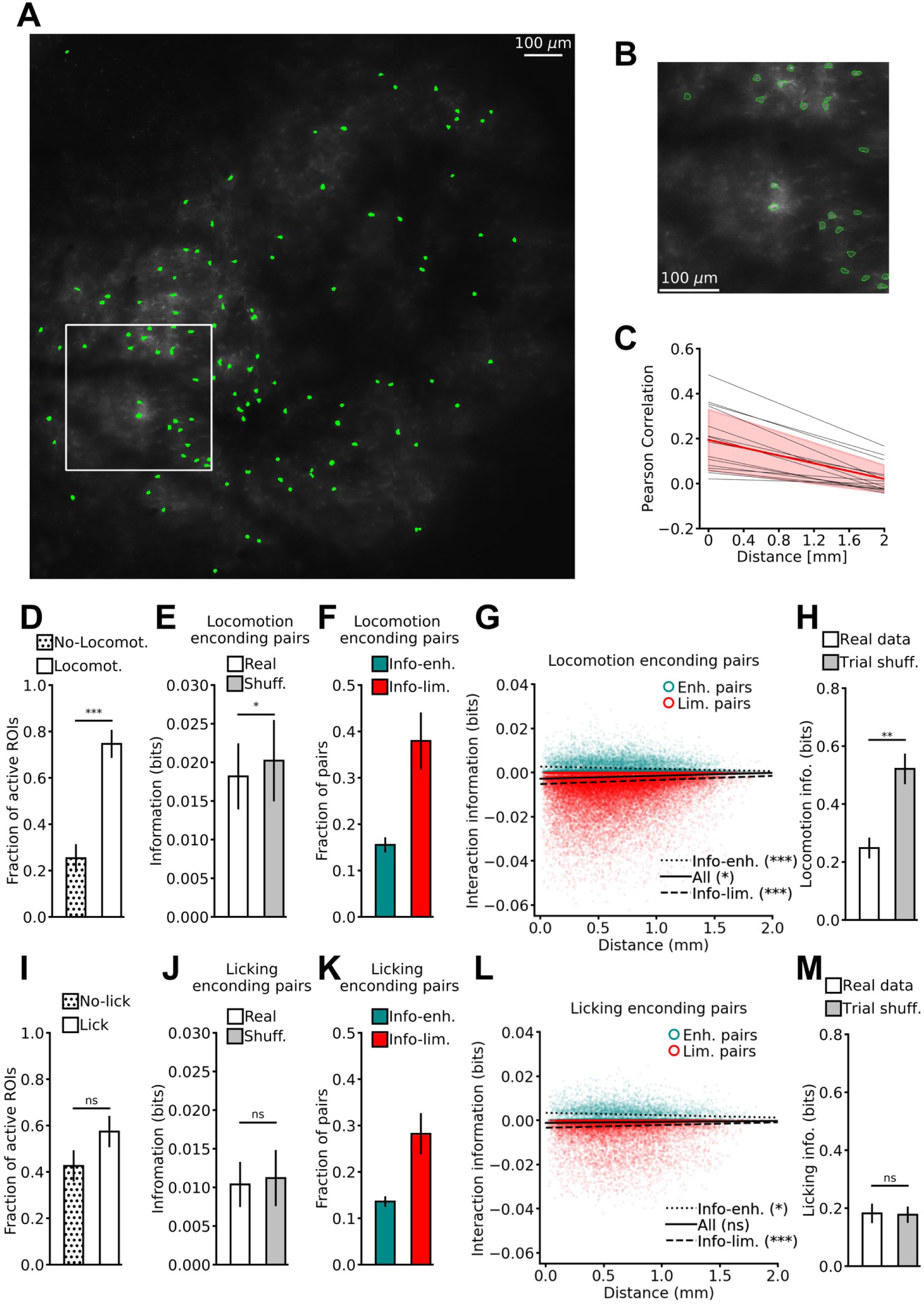
ASTRA enables fast segmentation of large-scale astrocytic networks imaged with two-photon mesoscopic microscopy. A) Representative image of somata segmentation (green) identified by ASTRA on cortical astrocytes expressing GCaMP6f and recorded using two-photon mesoscopic imaging (FOV dimension: 1.5 mm x 1.5 mm; FOV Id: 4). B) Zoom in of the region highlighted in A. On A-B image contrast has been adjusted to aid visualization. C) Pearson correlation of calcium signals for pairs of astrocytic somata as a function of pair-wise distance. Grey lines are linear fit of data from individual sessions, the red line is the mean ± std across 15 imaging sessions from 3 animals. Two-sided Wilcoxon rank-sum test. D) Fraction of active ROIs during animal locomotion (Run) and absence of locomotion (No-Run). Two-sided Wilcoxon signed rank test N = 13. E) Mutual information about animal locomotion carried by pairs of ROIs (I) compared with the sum of the information separately encoded by each member of the pair (ILIN) plus the signal similarity information component (Ish = ILIN+ ISS). Two-sided Wilcoxon signed rank test, N = 13. F) Fraction of information-enhancing (Info-enh.) and information-limiting (Info-lim.) pairs encoding locomotion information. N = 13. G) I - Ish values within pairs of somata as a function of pairwise distance. Information-enhancing pairs are reported in cyan and information-limiting pairs are reported in red. Linear regression fit: all pairs (solid line), information-enhancing pairs (dotted line), and information-limiting pairs (dashed line). Two-sided Wilcoxon signed rank test, N = 13. H) Information about locomotion behavior (Irun) decoded from astrocytic population vectors on real (white), and trial-shuffled (gray) data. Two-sided Wilcoxon signed rank test N = 13. I-M) Same as in D-H but for licking (lick) vs no liking (No-lick) behavior. I, two-sided Wilcoxon signed rank test N = 13; J, two-sided Wilcoxon signed rank test N = 8; K, N = 8; L, two-sided Wilcoxon rank-sum test, N = 8; M, Two-sided paired t-test., N=8. In (H-M) trial shuffling disrupted temporal coupling within astrocytic population vectors, while preserving single ROI activity patterns. In panels D, E, F, H, I, J, K, and M, data are represented as mean ± sem.

We next extracted the calcium fluorescent traces from each detected astrocytic soma and we used the extracted traces to characterize the network-level interactions of large-scale astrocytic populations. We first computed the Pearson correlation between the calcium activities of all pairs of somatic astrocytic ROIs. Activity correlations (Fig. 6C) were on average positive and larger than those typically observed when imaging the activity of neurons with calcium indicators, e.g., ^15, 49^. Correlation strength decreased as function of distance, but remained above zero even up to spatial distances of 2 mm, implying that astrocytes are functionally organized over mm-scale networks.

Activity correlations between neurons profoundly shape how neurons encode and transmit information at the level of large neural populations ^50–54^. However, little is known about how activity correlations of astrocytes influence the encoding of information about external variables in astrocytic networks. We investigated whether astrocytic activity correlations increased or decreased the information encoded by populations of astrocytes about two external variables: *i)* locomotion, i.e. whether or not the animal was running, and *ii)* licking, i.e., whether or not the mouse used its tongue to reach a water spout. For each cell pair, we computed the interaction information, quantifying how much correlations influence the information on a population code. This quantity is defined as the difference between the information about the external variables encoded in the “real data” activity of the pair, (Fig 6E,J) which contains contributions of both the tuning of the individual cells and of their correlations, and the “shuffled” information (Fig 6E,J) that the cell pair would carry if the cells have the same tuning to the stimuli as in the actual data but correlations are removed ^55, 56^. Positive (negative, respectively) interaction information values indicate that correlations enhanced (limited, respectively) the information encoded by the cell pair. For a given pair of cells, theoretical work, widely validated on neurons, demonstrated that positive activity correlations enhance or limit information, according to whether cells have similar or different tuning to the external variables, respectively ^50, 52, 57^. We found (Fig 6G,L) both information-limiting and information-enhancing correlations in astrocytic pairs, similarly to what previously reported for neurons ^52^. This could be explained by the same principles previously found in neurons. In fact, a large majority of astrocytes elevated their activity when animals ran (Fig. 6D), whereas comparable fractions of astrocytes elevated and decreased their activity when animals licked (Fig 6I). Thus, while most astrocyte pairs shared similar tuning to locomotion, a more balanced fraction of astrocytic pairs with similar *vs* different tuning to licking was present. As a result of the stronger homogeneity of tuning to locomotion, pairs of astrocytes had similar locomotion tuning and positive activity correlations. This led to a large majority of astrocytic pairs with correlations limiting locomotion information (Fig 6F-G). For licking, the greater diversity of tuning led to a more even distribution of positively-correlated pairs with either similar (information-limiting correlations) or dissimilar (information-enhancing correlations) licking tuning. Thus, because of the greater diversity of tuning for licking, the fraction of pairs with significantly information-enhancing and significantly information-limiting correlations was more balanced (Fig. 6K-L).

On average across all astrocytic pairs, there was a significant but moderate averaged information-limiting effect for locomotion information (Fig. 6E) and no effect for licking information (Fig. 6J). The average effect on pairwise information of correlations decreased as a function of distance for both information-enhancing and information-limiting pairs. This was because the correlations strength decreased, but remained sizeable, over distances > 1 mm (Fig. 6G, L).

Given that we obtained the first simultaneous recording of hundreds of astrocytes, we could evaluate how much information about licking and locomotion was encoded in large astrocytic populations, far beyond single cells and pairs, and how activity correlations shaped it. Studies with neurons showed large quantitative differences between encoding in small and large neural populations ^51, 52, 58^. We computed the decoded information about the animal’s locomotion from a Support Vector Machine (SVM) operating on population vectors of mesoscopic astrocytic ROIs. In the large population, we found a large information limiting effect for locomotion (Fig 6H, the total information in population activity was more than 2 times smaller than the information in the trial-shuffled responses). This effect emerged because the relatively small, yet predominant, pairwise information-limiting effects summed up in the large population (Fig. S13E). Conversely, when considering the information about licking encoded by the large-scale astrocytic population, we found that there was no effect of correlations on population information (Fig 6M). This result could be explained because the pairwise information-enhancing effects for licking were compensated by pairwise information-limiting effects (Fig. S13F).

We also considered the total amount of information carried by the mesoscopic astrocytic population. The population of all astrocytes in the mesoscopic FOV carried ∼ 0.2 bits of information about both locomotion (Fig. 6H) and licking (Fig. 6M), an increase of a factor of more than 10 with respect to the corresponding single cell values. Interestingly, the population of all astrocytes carried approximately the same amount of population information about licking and locomotion, despite the fact that the single-cell information values were almost twice as small for licking compared to locomotion (cf. Fig. 6H-M with Fig. 6E-J). This is because for licking the higher tuning diversity and the consequent lower information limiting effect made different cells less redundant and allowed licking information to build up more at the population level. Importantly, considering populations of astrocytes coming from more restricted spatial regions, such as those of size comparable to the FOV of traditional two-photon microscopes, would have given much lower information values (Fig. S13G-H), confirming the power of mesoscopic imaging for revealing large information-encoding network of astrocytes.

Taken together, these results demonstrate that information about external variables (locomotion and licking) is encoded across large-scale astrocytic networks spanning several mm of cortical tissue. These distributed networks of astrocytes are endowed with emergent properties due to their correlations, which are highly specific to the information content (that is, to the specific external variable being encoded).

### ASTRA processing time

We measured ASTRA processing time for the whole inference pipeline, repeating the analysis over 10 iterations. To allow for performance comparison across different configurations of hardware resources, we used a 26.3*10^3^ µm^2^ FOV and we artificially changed the t-series length from 300 to 700 frames. We compared three computational resource configurations: 4 CPUs, 20 CPUs, and 20 CPUs + 1 GPU (Fig. S14A). These configurations were chosen to compare the time needed to perform ASTRA analysis from laptop-like performance (i.e., 4 CPUs) to high-performance workstation equipped with computing accelerators (i.e., 20 CPUs and 20 CPUs + 1 GPU). We found the 4 CPUs configuration was the slowest configuration (12.33 ± 0.08 s for 700 frames, mean ± std) to detect and semantically segment astrocytic somata and processes. In contrast, the 20 CPUs configuration and the usage of a GPU accelerator reduced ASTRA processing time (7.27 ± 0.03 s for 700 frames with GPU *vs* 10.80 ± 0.09 s without GPU). The cross-correlation module was a significantly time-consuming block in the inference pipeline. We found that GPU computational power reduced the execution time of the cross-correlation computation (0.919 ± 0.004 s for 90 s t-series, mean ± std) compared to the 20 CPUs implementations (19.23 ± 0.09 s for 90 s t-series) and the 4 CPUs implementation (26.39 ± 0.04 s for 90 s t-series; Fig. S14B). We finally measured the processing time to detect and semantically segment somata and processes of astrocytes for large-scale mesoscopic and high sampling rate non-mesoscopic recordings using the 20 CPUs + 1 GPU configuration. We measured processing time of the inference pipeline for 10 iterations on FOV areas of 0.16 mm^2^ and 0.26 mm^2^ changing artificially the t-series length from 1000 to 5000 frames (Fig. S14C). ASTRA processing time was 22.1 ± 0.3 s for the 0.16 mm^2^ FOV and 25.7 ± 0.1 s for the 0.26 mm^2^ FOV both composed of 5000 frames.

These results demonstrate that ASTRA performed fast (few tens of s) semantic segmentation of astrocytes in two-photon t-series, speeding up and facilitating the analysis of astrocytic calcium signals in vivo.

## Discussion

Astrocytes, the main type of glial cells in the brain, display complex intracellular calcium dynamics ^4, 5, 46^ that can be captured with two-photon microscopy. These complex dynamics can be spatially localized in subcellular astrocytic ROIs, involve large portions of the astrocyte cell, and be coordinated across the astrocytic network ^20, 21, 23, 25^. Moreover, calcium signals in astrocytes encode information about synaptic function, circuit activity, and cognitive states ^2, 3, 59^. Having efficient tools for the analyses of these signals and the precise identification of astrocytic ROIs is thus of fundamental importance to study the physiology of astrocytic and neuronal circuits. To identify and segment astrocytes, manual annotation is still largely used. A problem with this approach is that it does not scale to large datasets and fields of view, requires significant amount of human training, and intrinsically suffers from human-dependent variability. State-of-the-art approaches to analytically segment astrocytic ROIs in two-photon imaging experiments as, for example, GECI-Quant (Srinivasan et al. 2015) and CHIPS (Stobart et al. 2018) provide semantic segmentation of astrocytes. These methods require careful setting of multiple user-defined parameters, which may vary with data acquisition conditions and with SNR, and require significant computation time. Moreover, CHIPS only segments active processes. The point discussed above limit the reliability and scalability of these approaches. Other analytical methods identify calcium events within and across astrocytes based on pixel-wise correlated dynamics ^14, 28–31, 60^. However, event-based approaches are computationally demanding, making it challenging to apply them to large datasets and fields of view. Moreover, event-based methods identify ROIs, but do not relate identified ROIs to the morphology of individual astrocytes. ASTRA enables the identification of astrocyte somata, processes, and domains, is scalable to large datasets and fields of view, addressing most limitations of current state-of-the-art methods.

### ASTRA achieves fast automated segmentation

ASTRA was designed to minimize computational time and its pipeline was massively parallelized on GPUs, enabling fast DNN training and inference steps. For example, ASTRA required few tens of seconds to process a t-series of average length and could segment a whole dataset of previously published data ^15^ in few minutes, reducing analysis time by more than a factor of 60 compared to current methods ^10, 19, 27^ and by almost three orders of magnitude compared to manual annotations. This feature of ASTRA was fundamental to enable scalability of ASTRA to large datasets, as for example the field-of-view of mesoscopic two-photon imaging (see discussion below) and may enable closed-loop experimental designs.

### ASTRA provides precise and reliable segmentation

ASTRA performance in astrocyte segmentation was similar to that of human annotators. We used three different annotators to manually identify and segment somata and processes in our datasets. We combined these annotations into a consensus annotation. Annotators showed large level of agreement in the segmentation of somata and lower level of agreement in the segmentation of processes in all the datasets, highlighting the variability of this manual task. Once trained on the consensus, ASTRA provided reliable and reproducible segmentation, avoiding human operator-dependent variability. Moreover, in this study we shared our imaging dataset, annotations, and codes. In future work, this open access repository may be enriched with additional segmentations by users from other laboratories, initializing the process of generating consensus annotations agreed across laboratories. Additionally, our datasets and annotations can be used as an online platform for benchmarking new computational algorithms for the analyses of astrocytic functional imaging recordings.

### ASTRA outperforms current methods and can be combined with event-triggered approaches for combined morpho-functional segmentation

We found that ASTRA performance was better than that of the multiple state-of-the-art tools for the analysis of astrocytic (e.g., GECI-Quant ^19^) and neural signals (e.g., CaImAn ^36^, UNet2DS ^39^, and STNeuroNet ^37^) that we tested. Compared to GECI-Quant, a main algorithm for the segmentation of astrocytic functional images, ASTRA did not require user intervention while the semi-automatic implementation of GECI-Quant required the user to define at least one specific intensity threshold for each astrocyte based on a temporal projection of the imaging session. This GECI-Quant feature resulted in a time-consuming procedure that limited the reproducibility of this approach and its scalability to large datasets. Conversely, ASTRA’s automatic pipeline used both spatial and temporal information to reliably generate a highly informative projection, which is semantically segmented by the specific design of the DNN. Several event-based methods have also been implemented to characterize the spatio-temporal patterns of astrocytic calcium activity ^28–31^. However, these algorithms require several user-defined parameters, which often depend on imaging conditions, SNR regimes, and fluorescent indicator. Moreover, the algorithmic complexity of these methods scale poorly with the size of the input sample (i.e. number of signal sources per sample, number of pixels per frames, and number of frames). Semantic segmentation performance of ASTRA was robust and reproducible and can thus be used to mitigate some of the limits of event based methods. ASTRA performance was stable over a wide range of PSNR situations and ASTRA performance was crucially aided by efficient feature engineering on two-photon images in the pre-processing step.

Importantly, ASTRA can be coupled with event-based segmentation methods, such as AQuA ^28^. Event-based segmentation approaches identify ROIs relying on the correlated temporal dynamics of calcium signals across individual pixels ^28–31^. ROIs identified with event-based methods, however, are not usually related to morphological features of astrocytes ^28–31^. To this end, we used ASTRA to obtain fast segmentation of individual astrocyte domains and then we seeded AquA using the domain ROIs identified by ASTRA. With this combined strategy, we enabled the extraction of morpho-functional information from individual astrocyte domain in population imaging recordings.

ASTRA outperformed multiple state-of-the-art methods for detection and segmentation of neurons in two-photon imaging recordings. We understand the lower performance of neuronal tools when applied to the analysis of astrocytes as arising from the major differences in morphological structure and in the timescales of calcium signals between neurons and astrocytes. Neuronal tools based on non-negative matrix factorization ^33, 36^ heavily rely on hypotheses of spatial contiguity and temporal activity organized over time scales of tens to hundreds of milliseconds. These hypotheses are ill suited to describe subcellular astrocytic calcium signals, which show heterogeneous spatial extents, diverse dynamic properties at the somatic and processes compartments, and slower timescales. Neuronal algorithms based on DNNs, such as Unet2DS ^39^ and STNeuroNet ^37^, albeit sharing architectural similarities with the DNN module of ASTRA, showed specific limitations. For example, Unet2DS allowed some degree of astrocytic somata detection. However, Unet2DS failed to perform semantic segmentation of astrocytic processes, especially in the absence of the pre-processing step aimed to enhance spatial and temporal features of astrocytic signals. Conversely, DNNs designed to extract activity localized at specific spatial footprints and temporal scales (such as STNneuroNet ^37^) do not fit the spatial and temporal features of astrocytic calcium signals. These considerations showcase some of the reasons underlying the better performance of ASTRA in comparison to the state-of-the-art methods for neuronal analysis that we tested, and highlight the necessity to develop algorithms and computational architectures specific for astrocytes, as done in this study.

### ASTRA generalized across acquisition conditions and fluorescence indicators

We tested ASTRA on four datasets that differed for acquisition parameters (galvanometric mirror-based scanning *vs* resonant mirror-base scanning; pixel size, 0.7-1.1 µm/pixel) and fluorophore type (GCaMP6f *vs* TdTomato). Importantly, it was possible to use ASTRA, endowed with its default weights pre-trained DNN on dataset-1, on never-before-seen data (datasets 3-4), achieving high detection and segmentation performance. Moreover, ASTRA performance in ROI detection and segmentation was comparable to that of human annotators for all datasets. These results suggest ASTRA is a flexible analytical tool that can be applied to heterogeneous two-photon imaging datasets of astrocytes. The high performance of ASTRA benefitted from transfer learning (as we started training from DNN weights pre-trained on a large dataset natural images) and from pre-processing specifically designed for astrocytes. However, while we used for training datasets that can be considered of reasonable size in the field of two-photon astrocytic recordings, the amount of data used for training was still limited with respect to those used to train DNNs on other tasks such as the identification of objects in natural images ^43, 61, 62^. As a result, it is possible that ASTRA may not work well under some conditions for never-seen-before data. To alleviate this concern, we included in ASTRA the possibility to further train its DNN on new annotated images that may better suit the setup at hand.

### Analysis of two-photon mesoscopic functional imaging of cortical astrocytes using ASTRA

Given its speed and performance, we used ASTRA for fast automated segmentation of large-scale mesoscopic imaging data comprising hundreds of astrocytes. This type of data is challenging for current analytical methods and is unpractical for manual annotation. Yet, mesoscopic two-photon imaging ^48^ has the potential to unravel the extent to which astrocyte interact and how they organize at the level of large networks. We here performed the first two-photon mesoscopic imaging of GCaMP6f-expressing astrocytes in awake head-restrained animals and then applied ASTRA to segment the mesoscopic acquisitions. This made it possible to obtain the first simultaneous calcium imaging analysis of networks of hundreds of astrocytes over the mesoscale. Using ASTRA, we found that calcium dynamics of distributed ensembles of astrocytes carried information about external variables, such as licking and locomotion, and that their calcium activity was correlated over large spatial scales. This finding suggests that astrocytes form extended information-bearing networks spanning several mm of cortical mantle. Moreover, we observed that these activity correlations had a major influence on the emergent properties of astrocytic population codes and that this influence strongly depended on the content of the information of the astrocytic population code. For locomotion, activity correlations had a profound information-limiting effects on the populations, because of the homogeneity of tuning of astrocytes. In contrast, activity correlations did not limit the population information when considering licking.

Thirty years of combined recording and analysis of neural populations contributed to lay down the foundations for the emergent principles of organization of neural population codes and their contribution to multiple brain functions ^50, 52^ . The results presented in this study suggest that coupling analysis methods such as ASTRA with tools of mesoscopic imaging can reveal as rich emergent properties in large astrocytic networks.

To conclude, we developed a novel DNN-based tool to achieve fast, precise, and automated semantic identification of ROIs in two-photon imaging experiments of large-scale astrocyte calcium signals. Our method enables automated astrocyte segmentation of mesoscopic two-photon imaging of astrocytes, revealing distinct behavioral-dependent population coding properties in mm-scale astrocytic network. Moreover, our shared dataset, annotation, and codes offers the field the possibility to achieve community-based consensus ground truth for astrocyte segmentation and a ready-to-use tool to benchmark new computational developments.

## Materials and Methods

### ASTRA algorithm

#### General information about the use of data and the pipeline

The workflow of ASTRA has two different pipelines: training and inference (Fig. 1A-B). Each pipeline analyzed a dedicated training or inference dataset, respectively. The training set was a dataset (for example, a subset of FOVs with annotated t-series) which was used to optimize the algorithm finding adequate pre-processing hyperparameters and DNN weights (see *Activity Map Generation Module, Local Activity Filtering Module, and Deep Neural Network Module*). The inference set was a dataset, completely separated from the training one (for example, another subset of FOVs), which was used to evaluate the performances of the algorithm.

Consensus annotated two-photon t-series recordings of astrocytes in the training set underwent a pre-processing procedure comprising Spatial sharpening, Putative bounding box extraction, and Local activity filtering (see Fig. S1C red flow chart path); preprocessed data were used to optimize the DNN weights (see Deep Neural Network Module). Analogously, t-series in the inference set underwent a similar pre-processing procedure, with the difference that data entered the pipeline without annotations (Fig. S1C blue flow chart path). The algorithm proceeded detecting putative cells which were then segmented by the DNN (Fig. S1B).

When we tested ASTRA on dataset-1 or dataset-2 (see *Generation of the two-photon imaging dataset in awake head-restrained mice*), we first trained the algorithm from scratch and then we tested it using leave-one-out cross-validation (the training set consisted of all but one t-series which was held out and tested as inference dataset). Results of these tests are reported as the averages across the leave-one-out replicates of training and inference.

We obtained the two sets of default weights distributed with the software training the DNN on the entire dataset-1 and dataset-2, respectively. When we performed inference on dataset-3 and dataset-4, we used the default weights obtained from the entire dataset-1.

The version of ASTRA released in this article can readily perform inference using the default values of pre-processing hyperparameters and DNN weights trained on dataset-1 or dataset-2. Both sets of parameters are distributed with the released software. New users can further optimize ASTRA by adding to the training pipeline their own t-series annotated with ImageJ (Fig. 1A, Fig. S12).

In the following sections, we provide detailed descriptions of the modules of ASTRA.

#### Spatial Sharpening Module

This module performed spatial sharpening and pixel intensity standardization on the median projection of a t-series. First, the module subtracted from each frame the 10^th^ percentile of the pixel intensities ^10^, then it computed the median projection on the entire t-series. Median projection’ pixels intensity was then standardized and rescaled as a 16-bit integer (i.e. within the interval [0; 2^16^]). Image contrast has been adjusted by using clipping limited adaptive histogram equalization (CLAHE, ^63^ OpenCV-python). Projections were then convolved with a sharpening kernel K.

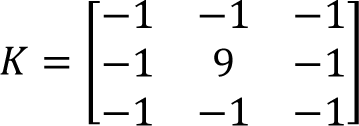

This set of operations condensed information about the spatial location of astrocytic signals collected over time into a single, highly informative, spatial map.

#### Activity Map Generation Module

This module detected regions in FOVs characterized by spatially-localized high fluorescence intensity (see below), generating a putative “activity map”.

As a first step, the input FOV was subsampled in overlapping patches (Fig. S1D), each subject to independent statistical analysis. Each patch was a 3D tensor in time and space in which the intensity value of each 3D voxel was considered an independent sample. For each time t, Voxels-vi,j(t), where i,j were indexes over the patch dimension, were binarized setting their value to 1 if their fluorescence intensity values were greater than the N-th percentile of the voxels intensity distribution within the patch or set to 0 otherwise. The N-th percentile was selected by optimization of the activity map generator performances on the training set (see below). Finally, a bi-dimensional (spatial) average projection of the binarized 3D tensors was generated reporting the fraction of time in which the voxels vi,j were classified to 1. In the areas of patch overlap, a bi-dimensional average projection for each pixel the spatial average was computed as the average value across patches. To provide biologically relevant constraints to this statistical filter, the number of astrocytes in each FOVs was estimated as the ratio of the FOVs surface with respect to the area of an astrocytic domain. Here, each astrocytic domain was approximated as a circle of surface π(d/2)^2^, where d was the characteristic diameter of an astrocytic domain (∼40 µm ^1, 45^). The estimated number of astrocytes represented a lower bound for the number of active zones; in fact, the number of identified clusters could be greater than the estimated number of astrocytes because of portions of astrocytic bodies visible in the FOV or blood vessels appearing as active areas. Finally, the algorithm identified all the spatially contiguous active clusters of pixels on the bi-dimensional (spatial) average projection of the binarized 3D tensors, selecting clusters with surface greater or equal than a putative somata (∼40 μm^2^). As a conservative initialization, pixels were considered active if their average projection value was greater than 0.6 (i.e. being classified as 1 on 60% of time frames). If the number of clusters identified was smaller than the estimated number of astrocytes the algorithm decreased by 0.03 the threshold for selecting active pixels. For each new threshold the putative somata surface area was decreased iteratively by 4 µm^2^ starting from 40 μm^2^ to 20 μm^2^. This tuning process stopped when the number of identified clusters is equal or greater than the estimated number of astrocytes the algorithm. This procedure aimed to minimize the difference between the number of detected active regions and the theoretical astrocyte number.

The N-th percentile used to binarize the 3D tensor was tuned optimizing the performances of the activity map generator module on the training set. Performances of this module were evaluated computing the F1-score value between consensus somata annotations (see Manual Dataset Annotation section) and active zones identified in each FOV of the training-set. The performance was computed for a set of percentiles (30, 40, 50, 60, 70, 80, 90) and the one which maximized the F1-score was selected.

#### Putative Bounding Boxes Extraction Module

This module computed centroids of active zones detected by the activity map generation module and generated bounding boxes (BBs) surrounding them. BBs were ∼55 μm high and wide. These BBs were used to extract from t-series and spatial sharpened maps respectively (putative) single-cell spatial map and (putative) single-cell recordings, respectively.

#### Local Activity Filtering Module

This module performed a fine local time filtering on single-cell recordings. The module computed the 90^th^ percentile of the pixels intensity distribution and used it as a threshold to binarize the tensor. Then the module selected the pixels which were set to 1 for at least the α1% of the frames. This binarization procedure was repeated setting to zero the previously selected pixels from the starting distribution. Hence, all the pixels which were set to 1 for at least α2% of the frames were selected. The binary mask obtained from the summation of the two previous set of pixels was used to generate a binary map (1 foreground and 0 background) to filter background regions and eventual artifacts generated by spatial sharpening from spatial single-cell spatial maps.

The thresholds α1 and α2 used in this module were tuned on the training set. The module explored a set of α1 and α2 couples ([0.3,0.15], [0.25,0.1], [0.2,0.07], [0.15,0.05]) and computed the fraction of pixels belonging to consensus annotation removed by the filter. Finally, the module selected the couple (α1, α2) with the highest threshold values that removes less than 5% of consensus pixels for both soma and processes (i.e. α1=0.25 and α2=0.10 for dataset-1). We provided these values as defaults for new users who will decide to run inference with ASTRA with default hyperparameters and weights without further optimizing it using new annotated data.

#### Deep Neural Network Module

Our design of this convolutional Deep Neural Network (DNN) started from a U-net ^39^ architecture with an encoder part (the descending part of the U shape in Fig. S1E) that analyzed the input image and a decoder (the ascending part of the U shape in Fig. S1E)) that took the information from the encoder and up-sampled it to classify the pixels of the input image. The first two blocks of the encoder (L1 and L2 of the left part in Fig. S1E) were two basic U-net blocks that analyzed input images using convolutional filters. We then nested three pretrained Inception-Resnet-v2 modules ^64^ in our network (L3 to L5 levels in the left part in Fig. S1E) changing its encoder backbone ^65, 66^.

The decoder part of the U-Net (right part of Fig. S1E) implemented in 5 levels (L5 to L1) an up-sampling strategy that was a fundamental transformation operation to obtain a pixel-level prediction of the class with which each pixel should be labeled. In the Decoder part, we adopted Dense Up-sampling Convolution (DUC) to reduce the decoder number of weights ^67^. The DNN outputs consisted in a 3D tensor whose dimensions were: input height, input width and 3 channels corresponding to the probability of the pixel to belong to somata, processes, or background classes. Each pixel was finally assigned to the class with the highest probability.

During training of all the layers of both the encoder and decoder parts, we used data augmentation techniques to limit algorithm overfitting problems caused by the relatively small size of the dataset. During training, we used standard transformations ^68^ of input images: rotation by 90°, 180°, 270°, Gaussian blurring with a 3×3 pixels kernel and σ=3, Gaussian noise sampling values from a Gaussian distribution with μ = 0 and σ = 0.3, salt and pepper noise on 4% of pixels, scaling of image size by factor 1.4, 0.9 and 0.8, horizontal and vertical flipping, pixels intensity scaling by factors 3 and 0.5. Moreover, we used morphological transformations that altered the spatial structure of input images: elastic (Ronneberger, Fischer, and Brox 2015), barrel, and pincushion.

We combined a Binary-Cross-Entropy (BCE) loss with soft Dice loss (Milletari, Navab, and Ahmadi 2016); BCE was applied to all the three classes soma, process, and background. Soft Dice has been applied only on soma and processes:

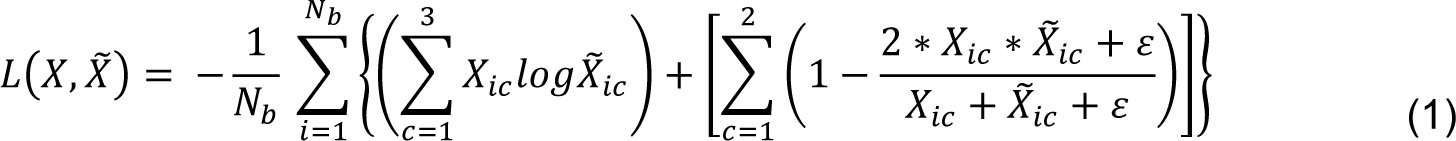

where X and 𝑋̃ represented user defined mask tensor and prediction mask tensor, ε = 0.5 was constant that preserved the numerical stability, 𝑁_𝑏_ was the batch size and c=1,2,3 was the class index for: processes, soma, and background. The role of DNN was to assign small regions to the correct class, hence, the soft dice loss represented a proper metric to measure area overlapping accuracy. We trained the DNN using Adam Optimizer ^69^ and starting learning rate lr (see SI Tab. S7). The number of training epochs was N = N_1_+N_2_. In the first N_1_ epochs of training, the weights of pretrained blocks (Reduction Blocks and IncecptionResNet Blocks) were not updated. During the remaining N2 epochs, we performed a fine tuning of the entire net weights. All the filters trained since the first epochs were initialized as described in ^65^. Training details are reported in SI Tab. S7.

#### Cross Correlation Module

This module analyzed fluorescence intensity dynamics of pixels within the putative domain surrounding the semantically segmented astrocytic soma and processes ROIs (i.e. a circular region of radius ∼40 μm). We referred to the intensity fluctuations in time of each pixel as 𝑠(𝑡). This module was composed of two blocks – cross-correlation computation and threshold optimization – that were executed iteratively. The cross-correlation computation block classified a set of input 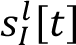 for l =1, …, L as correlated to a set of reference 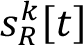 or k =1, …, K given a threshold th_cc_; where L was the number of inputs and K is the number of references. The block computed the normalized cross-correlation between each (l,k) couple (Eq. 2) and selected its maximum value (Eq.3). Then, the cross-correlation matrix M_cc_ is defined as in eq. 4.

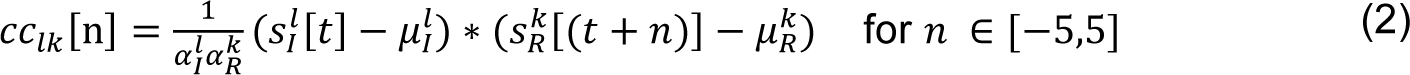

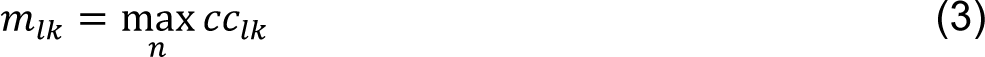

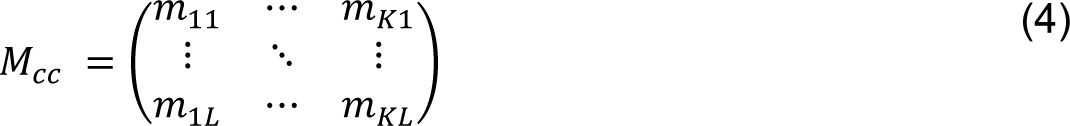

M_cc_ was then binarized selecting only values greater than threshold th_cc_. 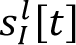 was classified cross-correlated if at least one element in the l^th^-row of M_cc_ was equal to one.

The threshold optimization block selected an optimal threshold using an iterative approach. A set of 250 pixels was sampled outside astrocyte domains in each FOV and their 𝑠_𝑃_[𝑡] were collected. This set represented a proxy over which we could compute the number of false positive selections obtained from the cross-correlation computation block using as a reference set the 𝑠_𝑅_ [𝑡] extracted from ROIs pixels. Since pixels were sampled outside astrocytic domains, these pixels could not belong to any subcellular region of the astrocytes detected in the FOV. For this reason, we assumed that the sampled pixels 𝑠_𝑃_ [𝑡] were independent from the ones of the semantically segmented ROIs pixels. In the threshold optimization block, for each threshold value in the range 0.60 to 0.95 with minimum spacing between values of 0.05, we computed the number of false positive pixels. Then, this block selected as optimal threshold thOp the smallest threshold value with average false positive percentage error less than 5 %. Finally, for each detected cell the cross-correlation module collected all the pixel 𝑠_𝐼_ ⌈𝑡⌉ in the circular region that surrounded it and all the ROIs pixels 𝑠_𝑅_ [𝑡]; then it applied the cross-correlation computation block on these two sets of 𝑠[𝑡] using thOp as threshold.

### Workflow pipelines

#### Training Pipeline

The training-phase was organized as a series of steps that end with the DNN training as shown in Fig. S1A. First, the spatial sharpening module was applied to the training FOVs generating spatial sharpened maps. Since cells in the training set were already segmented in the consensus segmentation, the Putative Bounding Boxes Extraction module generated the BBs using the somata annotated in the consensus segmentation as input. The Putative Bounding Box Extraction module generated a set of single cell spatial maps and a set of single-cell recordings. The Local Activity Filtering module analyzed the single-cell recordings obtaining binary masks of foreground/background pixels. Finally, single cell images extracted from the spatial maps were filtered with these binary masks. This filter further denoised and enhanced the so-obtained single cell spatial maps. This pipeline ended after the training of the DNN with the so obtained single cell filtered spatial map images.

#### Inference Pipeline

The inference-phase started with the pre-processing which generated a set of putative filtered single-cell maps from the inference set, as shown in Fig. S1B. The pre-processing was organized in several steps where Spatial Sharpening, Activity Map Generation, Putative Bounding Boxes Extraction, and Activity Filtering modules are applied. For each FOV, the spatial map and the activity map were generated by the Spatial Sharpening module and by the Activity Map Generation module, respectively (Fig. S2A). Then, the Putative Bounding Boxes Extraction module extracted the putative single-cell spatial maps and the putative single-cell recordings. Finally, the Activity Filtering module analyzed single-cell recordings and identified background zones. These zones were filtered from the single-cell spatial map (Fig. S2B). Subsequently, the filtered single-cell spatial maps were used to reconstruct a spatial map of the entire FOV where all the background parts were filtered. The DNN analyzed the filtered single-cell spatial maps and the segmentations of the DNN were placed at the correct location within the FOV using the BB coordinates. Altogether these segmentations constituted the semantic segmentation of the entire FOV. Then, the DNN analyzed the FOV filtered spatial map providing for each cell the probability of being a true-or a false-positive. Cell probability was computed as the mean probability of pixels inside somata ROIs of being classified as soma-type pixels by the DNN. Cells with probability smaller than 0.9 were filtered from the FOV segmentation results. The segmented regions obtained were spatially filtered including only cells with identified soma area greater than 0.9*Amin and smaller than 1.1*Amax, where Amin was the smallest somata area measured in the training dataset whereas Amax was the greatest somata area measured. Finally, identified processes were filtered if not spatially connected to an identified soma. If needed, users can then proceed to subcellular parcellation of the segmented processes setting a suitable surface value to split process-ROIs in “mini-ROIs”. The last step consisted in the refinement of the ROIs so obtained using the cross-correlation module. In fact, it identified regions where calcium signals were cross-correlated with the semantically segmented ROIs signals in the FOV.

### Detection and Segmentation Metrics

We evaluated the detection performances of our algorithm by comparing ASTRA somata segmentations with the manual consensus labels, as described in ^36, 37^. We quantified three somata detection metrics: recall, precision, and F1 score, defined as follows:

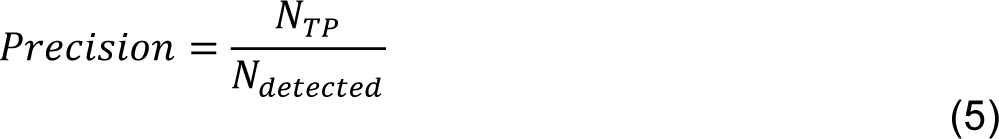

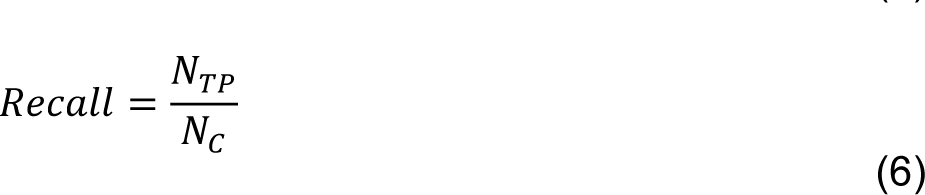

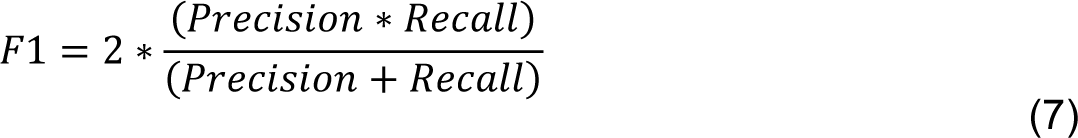

We defined these quantities as follows: number of manually labelled somata (consensus somata, 𝑁_𝐶_), number of true positive somata (𝑁_𝑇𝑃_) and number of somata detected (𝑁_𝑑𝑒𝑡𝑒𝑐𝑡𝑒𝑑_) ^36, 37^. We matched masks between the consensus labels and the detected masks using the Intersection-over-Union (IoU) metric along with the Hungarian algorithm ^70^. We computed the IoU metrics for 2 binary masks 𝑀_1_ and 𝑀_2_ as follows:

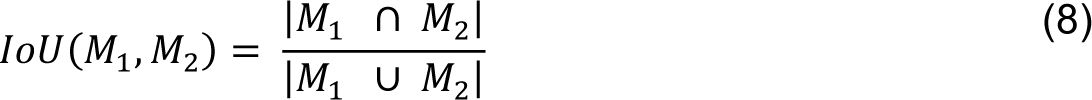

Then we computed the distance matrix between any pair of masks in GT manual annotations set and in ASTRA annotations set as described in ^36, 37^. Each element of this matrix has been computed as follows:

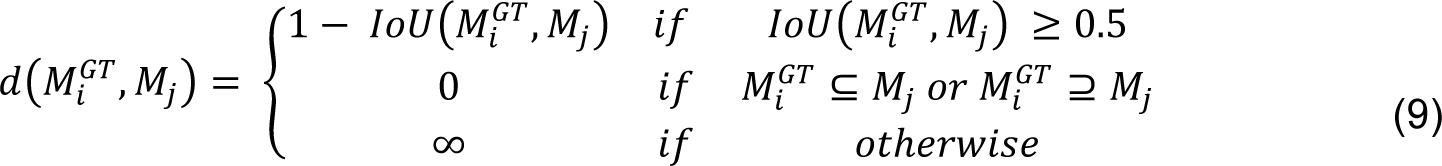

A distance of infinity corresponded to non-matching masks due to their small IoU score. Finally, we solved the matching problem applying the Hungarian algorithm to the distance matrix. The number of matched masks corresponded to 𝑁_𝑇𝑃_.

Segmentation scores have been computed at the pixel level to quantify how complex structures like processes were segmented by ASTRA. For each FOV, we computed the segmentation score considering only the detected cell; when no detected cells were available in a FOV the segmentation score was discarded. The segmentation score was quantified by three metrics: recall, precision, and F1 score, defined in eq.5, eq.6, eq.7. We defined 𝑁_𝐶_ number of manually labelled pixels, 𝑁_𝑇𝑃_ number of true positive pixels in the ASTRA segmentation and 𝑁_𝑑𝑒𝑡𝑒𝑐𝑡𝑒𝑑_ the number of pixels segmented by ASTRA. We computed the F1-score for both somata and processes pixel-classes.

### Cross-Correlation Error evaluation

Error estimation for the cross-correlation module has been performed computing the number of pixels outside astrocyte domains that were cross-correlated with the consensus ROIs pixels in each FOV.

For each cell in the FOVs, we sampled 1000 pixels outside the astrocytes’ domain avoiding pixels which were used to tune the cross-correlation threshold. This set was fundamental to compute the number of false positive selections for each astrocyte. Domains were estimated as a circular region of radius ∼40 µm surrounding each cell in FOVs. Then, we computed the number of false positive pixel per FOV_i_ as:

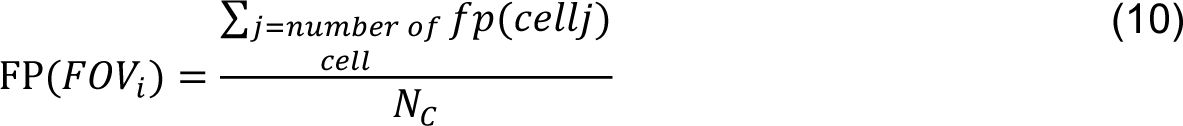

Where fp was the number of false positive pixels selected for cell_j_ and N_C_ was the number of cells in FOV_i_.

### Implementation of other state-of-the-art algorithms used for comparison

#### UNet2DS implementation

We segmented dataset-1 with UNet2DS software (https://github.com/alexklibisz/deep-calcium) validating its performance using leave-one-out cross validation strategy. For dataset-1, we used the same training procedure outlined in ^35^. We used 50 epochs with 100 training iterations in each epoch using sixteen randomly cropped 128×128 pixels regions from the mean image, utilizing the dice-loss and the Adam optimizer. We monitored the 𝐹1 score on a validation set, which was selected from the training set (5% of the training set) to ensure the network was not overfitting.

#### STNeuronet implementation

We segmented dataset-1 with STNeuronet ^37^, validating its performance using a leave-one-out cross validation strategy. We preprocessed our data as described in ^37^ and we adapted the consensus annotation (see *Consensus annotation*) to identify active somata of astrocyte in each frame of our data (https://github.com/soltanianzadeh/STNeuroNet, prepareTemporalMask.m). In the training dataset, somata were classified as active/inactive analyzing Δf/f_0_ traces extracted using the procedure described in ^15^ to detect statistically significant calcium events. For each FOV, we generated the training set cropping 120×144×144 voxels surrounding each somata in the consensus annotation. Then, we trained the net for 10000 epochs with leaning rate 0.5*10^-4 and batch size of 3. The loss function always converged to a plateau within 10000 epochs with these training parameters. Then, we used the same training procedure outlined by ^37^.

#### CaImAn implementation

We segmented dataset-1 with CaImAn ^36^, validating its performance against the consensus annotations. CaImAn hyperparameters were set according to astrocytic somata morphology ^1, 45^ and signal dynamics. We used patch_ size = [80, 80] and overlap = [20, 20] for dataset-1. Components to be found was set to K = 1 since in these patches there was at least 1 astrocytic somata. Decay time was 1.5 s and we set merging threshold equal to 0.6 in each test. Other parameters were set to default settings.

#### GECI-Quant

To perform semi-automatic semantic segmentation with GECI-Quant, annotator-1 followed the procedure described in ^19^. Briefly, for each FOV in dataset-1, the annotator selected two regions of interest for every astrocyte corresponding to soma and astrocytic domain, respectively. Then, annotator-1 manually selected an intensity threshold for each region of interest following the procedure outlined in ^19^. GECI-Quant segmentations were used to compute the performance.

### Reconstruction of astrocytic morphology from the spatio-temporal map of AQuA

Starting from the spatio-temporal map of calcium events resulted from AQuA ^28^, we reconstructed astrocytic morphology since a subset of pixels classified as events should in principle belong to astrocytic somata and processes. For each astrocyte detected in the consensus annotation, we ran AQuA in circular regions of radius ∼40 µm surrounding these cells, thus limiting the analysis to the putative astrocytic domain. Using putative calcium events detected by AQuA, we selected the pixels belonging to a minimum number of events. For each astrocyte the minimum number of events was tuned as the value that maximize F1-score between the selected set of pixels and the consensus annotation (using the union of somata and processes annotations). Hence, we computed precision, recall and F1-score between the best reconstruction and the consensus annotation. This strategy provided the F1-score upper bound for the reconstruction astrocytic morphology using AQuA.

### PSNR evaluation

We evaluated the peak signal-to-noise ratio (PSNR) of the FOV_i_ containing N astrocytic ROIs as:

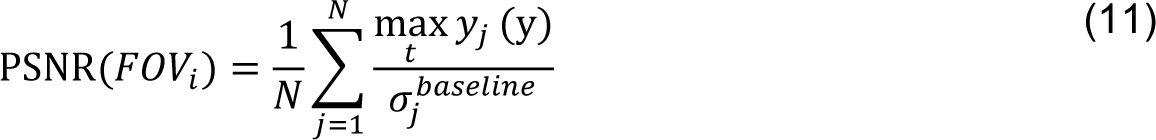

where y_j_ (t) was the mean fluorescence signal in astrocyte ROI_j_ and 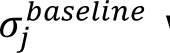 was the standard deviation of the baseline distribution of fluorescence values in astrocytic ROI_j_. To compute the baseline distribution of each astrocyte, we considered only pixels inside the astrocyte domain (circular area of radius ∼40 μm). The values of these pixels across time formed the full fluorescence distribution. The baseline distribution consisted of all the fluorescence value smaller than the 80^th^ percentile of the full fluorescence distribution.

### Manual dataset annotation

Two-photon t-series were motion corrected with a custom Python implementation of phase correlation correction algorithm ^71^. Motion-corrected t-series were pre-processed with the spatial sharpening module (see above). The consensus generation process included 2 steps. In the first step, three expert annotators independently labeled the datasets using the freehand and ROI Manager tools of Fiji ^72^ according to the following rules: i) annotators used the t-series to detect visible astrocytic somata; ii) spatial maps were used to select and to label ROIs, identifying visible astrocytic somata and processes; iii) annotators sequentially added ROIs, defining the contours of the optically resolved proximal processes displaying active calcium dynamics and presumably belonging to the same astrocyte.

In the second step, annotators solved inconsistencies in their annotations reaching a consensus ^36^ as follows: i) annotations of the three annotators were combined in overlapping masks (Fig. S3) to highlight discrepancies among annotators; ii) each soma or process identified by 3 annotators was included in the consensus; iii) each soma or process identified by < 3 annotators was included in the consensus only after an ad-hoc review, where the annotators judged looking at both the preprocessed spatial maps and motion corrected t-series.

### Extraction of calcium event traces

For each ROI, we computed fluorescence signals as in eq. 12.

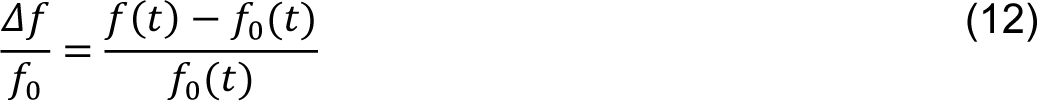

Where f(t) was the average fluorescence signal of a given ROI at time t and f_0_(t) was the baseline fluorescence. f_0_(t) was computed as the 20^th^ percentile of the fluorescence intensity in a 30 s rolling window centered in t. Then, we generated the calcium event traces of astrocytes following the procedure described in ^15^. For each *Δf/f_0_* trace, the standard deviation σ_1_ of the whole signal was computed. Values above and below the interval ± σ1 were removed from the trace and the standard deviation σ_2_ of the filtered trace was computed. Finally, fluorescence transients were identified on the original trace as events if: i) fluorescence values were above 2σ_2_; and ii) fluorescence values returned within the ± σ_2_ interval in more than 0.5 s. Hence, we generated the calcium event trace setting all fluorescence values in *Δf/f_0_* outside of those belonging to positive events to 0.

### Mutual information analysis of individual astrocytic soma and of pairs

For experiments in which we recorded astrocytic calcium activity using mesoscopic two-photon microscopy, we computed locomotion information (information about whether the animal was running or was still) and licking information (information about whether the animal was or was not licking the waterspout) carried by the calcium signals of either a single astrocytic soma or jointly by a pair of simultaneously recorded astrocytic somata.

Mutual information between the considered behavioral variable, ***S*** (describing licking or locomotion), and the astrocytic activity, ***R***, was computed as:

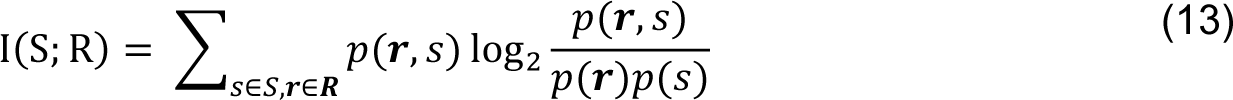

where1 *p(**r**,s)* is the joint probability of observing at the same time a value ***s*** and ***r*** for the behavioral and the astrocytic activity variable, and *p(s)* and *p(**r**)* are the marginal probabilities, respectively. The calcium activity of each individual soma was discretized in two equally spaced bins. For single soma information, ***r*** was a one-dimensional array reporting the discretized activity of the considered cell, and for pairs it was the two-dimensional array containing the discretized activity of the two somata. The behavioral variable locomotion was computed as a binary array, set to 1 for epochs of motion (mouse speed ≥ 0.5 cm/s) and set to 0 for epochs of immobility (mouse speed < 0.5 cm/s). The behavioral variable licking was a binary variable, in which contacts of the mouse’s tongue to the capacitive waterspout were encoded as 1 while licking was set to 0 otherwise. When considering pairs of ROIs, we further performed information breakdown analysis ^56, 73^, decomposing mutual information carried by a pair of ROIs, I(S;R), into four terms: 1) ILIN the mutual information linear term; 2) ISS the signal similarity term; 3) ICI the stimulus independent correlation term; 4) ICD the stimulus dependent correlation term.

We computed the null distribution for each pair of ROIs to evaluate if information in the two ROIs of the pair is information-enhancing or information-limiting. We generated n = 100 random shuffling of the behavioral variable label of the data, which destroyed the relationship between the behavioral variable and the calcium response. From the shuffled data, we computed the distribution of Ish = I – I_LIN_-I_ss_. A pair was classified information-enhancing or information-limiting if its real Ish value was >the 95^th^ percentile or < 5^th^ percentile of the shuffled distribution, respectively.

To correct the mutual information bias caused by limited sampling of astrocytic responses, we used the quadratic-extrapolation bias correction ^73, 74^.

### Mutual information computed from large astrocytic populations from the confusion matrix of an SVM decoder

To compute information for large populations of astrocytic ROIs, we computed mutual information with an intermediate decoding step ^75^, because we could not extend the direct information calculation of the previous sections to large populations due to sampling problems ^76^. We trained a support vector machine (SVM) ^77^ with Gaussian kernel to classify the state of either one of two behavioral variables (S) according to a single-trial population vector made combining calcium signals of all individual astrocytic ROIs within the FOV (either using all of them, or only part of it, see main text). Behavioral variables classified were locomotion and licking. Locomotion was defined as vector of binary values in which 1 or 0 indicated epochs of motion (mouse speed ≥ 0.5 cm/s) or epochs of immobility (mouse speed < 0.5 cm/s). Licking was defined as a binary vector: 1 when contacts of the mouse’s tongue to the capacitive waterspout were done and 0 otherwise. For each experimental sessions, the dataset was composed by Z_exp_ observations (X_j_,s_j_) with j = 1,…, Z_exp_. X_j_ is the n-dimensional array of the calcium activity of the N ROIs in the session, whereas s_j_ corresponded to either running/still or licking/not licking behavioral variables. Calcium observations X_j_ were used to predict s_j_ variables using a support vector machine with Gaussian kernel. We trained and tested the SVM using 5-fold cross-validation procedure independently on each experimental session. During each iteration of the cross-validation, optimal hyperparameters were selected performing 5-fold cross-validation on each fold training set.

Predictions of the decoder for each of the 5-folds used as test were then collected to compute the overall mutual information between the predicted behavioral variable S_p_ and the real value S. Mutual information I(S; 𝑆_𝑝_) was defined as the information in the confusion matrix:

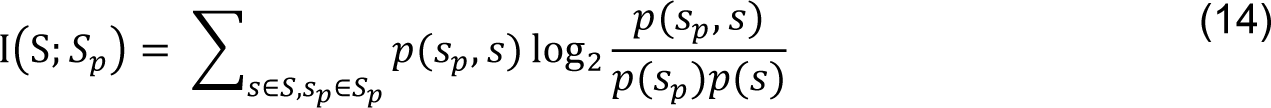

where 𝑝(𝑠_𝑝_, 𝑠) was the confusion matrix, that is the probability of observing a given value s of the behavioral variable and of predicting is as 𝑠_𝑝_, and 𝑝(𝑠) and 𝑝(𝑠_𝑝_) are the marginal probabilities, respectively. To assess if the correlations among astrocytes increased the amount of information related to a behavioral variable, we disrupted correlations by randomly shuffling, separately for each ROI, the order of trials with the same behavioral variable identity. We performed 100 trial shuffling. We then used the distribution of I(S; 𝑆_𝑝_) values on trial shuffled data to compute the trial shuffling information as the mean value of this distribution.

### Animals

All experiments involving animals were approved by the National Council on Animal Care of the Italian Ministry of Health and carried out in accordance with the guidelines established by the European Communities Council Directive authorization (61/2019-PR). All data were collected from male C57BL/6J mice (Charles River, Calco, Italy). From postnatal days 30, animals were separated from the original cage and housed in groups of up to five littermates per cage with ad libitum access to food and water in a 12-hours light-dark cycle. All the preparative and experimental procedures were conducted on animals older than 10 weeks.

### AAV injection and chronic hippocampal window surgery

Animals were anesthetized with 2% isoflurane 0.8 % oxygen, placed into a stereotaxic apparatus (Stoelting Co, Wood Dale, IL), and maintained on a warm platform at 37°C for the whole duration of the anesthesia. Before surgery, a bolus of Dexamethasone (Dexadreson, 4 gr/kg) was injected in the animal’s hamstring. After scalp incision, a 0.5 mm craniotomy was drilled on the right hemisphere (1.75 mm posterior, 1.35 mm lateral to bregma), the AAV-loaded micropipette was lowered into the hippocampal CA1 region (1.40 mm deep to bregma). 0.8 μl of AAV solution containing pZac2.1 gfaABC1D-cyto-GCaMP6f (Addgene viral prep # 52925-AAV5) was injected at 100 nL/min by means of a hydraulic injection apparatus driven by a syringe pump (UltraMicroPump, WPI, Sarasota, FL). Following the viral injection, a stainless-steel screw was implanted on the cranium of the left hemisphere and a chronic hippocampal window was implanted on the contralateral hemisphere similarly to ^15, 78, 79^. In brief, a trephine drill was used to open a 3 mm craniotomy centered at coordinates 2.00 mm posterior and 1.80 mm lateral to bregma. The dura was removed using fine forceps and the cortical tissue overlaying the hippocampus slowly aspirated using a blunt needle coupled to a vacuum pump. During aspiration the exposed tissue was continuously irrigated with normal HEPES-buffered artificial cerebrospinal fluid (ACSF). Aspiration was stopped once the thin fibers of the external capsule were exposed. An optical window was fitted to the craniotomy in contact to the external capsule and a thin layer of silicone elastomer (Kwik-Sil, World Precision Instruments) was used to surround the interface between the brain tissue and the steel surface of the optical window. The optical window was composed of a thin-walled stainless-steel cannula segment (OD, 3 mm; ID, 2.77 mm; height, 1.50 - 1.60 mm) and a 3.00 mm diameter round coverslip, which was attached to one end of the cannula using UV curable optical epoxy. Sharp edges and bonding residues were smoothed using a diamond coated burr. A custom stainless-steel head-plate was attached to the skull using epoxy glue. The components described above were finally fixed in place using black dental cement and the scalp incision was sutured to adhere to the implant. All the animals received an intraperitoneal bolus of antibiotic (BAYTRIL, Bayer, Germany) to prevent postsurgical infections.

### Generation of the two-photon imaging dataset in awake head-restrained mice

The optical setup for two-photon imaging was composed of a pulsed laser source (Chameleon Ultra, 80 MHz repetition rate tuned at 920 nm, Coherent) and Bruker Ultima Investigator equipped with 6 mm raster scanning galvanometers, movable objective mount, 16x/0.8 NA objective (CFI75 LWD 16X W, Nikon, Milan), and multi-alkali photomultiplier tubes. Laser beam intensity was adjusted using a Pockel cell (Conoptics Inc, Danbury). Laser beam power at the objective outlet was 90-110 mW. GCaMP6f or TdTomato emission signal was collected by the photomultipliers after band-pass filtering (525/70 nm) and digitalized in 12 bits. Imaging sessions were conducted in raster scanning mode. t-series were motion corrected using an open-source implementation of up-sampled phase cross-correlation ^71, 80^ and the t-series median projection was used as reference frame. One or two weeks after surgery the animals were handled by the operator for a minimum of two days and habituated to the imaging setup. Starting from the second session, the animals were head-restrained for a progressively increasing amount of time, reaching 1 hour in approximately one week. Mice were free to run on a custom 3D printed wheel. Experimental sessions lasted approximately one hour. After each session, animals were returned to their home cages.

#### Dataset-1

This dataset was composed of 24 t-series of hippocampal astrocytes expressing GCaMP6f, recorded in head-fixed mice running on a wheel. This dataset was composed of 15 t-series with 550 frames and 9 FOVs with 750 frames (image size, 256 pixels x 256 pixels, 0.634 μm/pixel. T-series were acquired at 3 Hz.

#### Dataset-2

This dataset was composed of 10 t-series of hippocampal astrocytes expressing TdTomato, recorded in head-fixed mice running on a wheel. 4 t-series were recorded with galvanometric mirror scanning and 4 t-series were recorded with resonant mirror scanning. Image dimensions were 512 pixels x 512 pixels, 1.057 μm/pixel. t-series recorded with galvanometric mirror scanning comprised 250 frames. t-series recorded with resonant mirror scanning comprised 1200 frames (one t-series), 5500 frames (two t-series), and 9000 frames (one t-series). t-series recorded with galvanometric mirror scanning were recorded at [0.8-1 Hz], whereas t-series recorded with resonant mirror scanning were recorded at 30 Hz.

#### Dataset-3

This dataset was composed of 7 t-series of hippocampal astrocytes expressing TdTomato indicator recorded in awake mice running on a wheel. t-series length was: 5500 frames (5 t-series), 4500 frames (one t-series), and 9000 frames (one t-series). Image dimensions: 512 pixels x 512 pixels, 0.79 μm/pixel. t-series were acquired at 30 Hz.

#### Dataset-4

This dataset was composed of 10 t-series of hippocampal astrocytes expressing GCaMP6f, recorded in head-fixed mice running on a wheel. The dataset comprised 10 t-series. t-series length, 9000 frames; image dimension, 512 pixels x 512 pixels; pixel dimension, 0.793 μm/pixel; acquisition frequency, 30 Hz.

#### Simulated datasets

We generated 4 artificial datasets with increased noise levels using dataset-1. To this aim, we first estimated the standard deviation, σ, for each pixel in the FOVs. We then computed a novel temporal intensity trace adding zero mean gaussian noise with β*σ standard deviation to the recorded raw trace. The noise scaling factor, β, was 0.5, 1, 1.5, 2 for the four artificial datasets, respectively.

We also generated 2 datasets with reduced background pixels intensity. For each t-series, we defined as background all the pixels outside the consensus annotations, and we scaled background pixel intensity by a factor λ, (λ values, 0.75 and 0.5, respectively).

### AAV injection and chronic optical window surgery for mesoscopic imaging of cortical astrocytes

Animals were anesthetized with 2% isoflurane 0.8 % oxygen, placed into a stereotaxic apparatus (Stoelting Co, Wood Dale, IL), and maintained on a warm platform at 37°C for the whole duration of the anesthesia. Before surgery, a bolus of Dexamethasone (Dexadreson, 4 gr/kg) was injected in the animal’s hamstring. After scalp incision, two small craniotomies were drilled on the right hemisphere (craniotomy 1: 1.25 mm posterior, 2.00 mm lateral to bregma; craniotomy 2: 1.75 mm posterior, 1.60 mm lateral to bregma). A micropipette loaded with AAV solution was lowered 300 µm below pial surface into the cortical parenchyma. 0.4 μl of AAV solution containing pZac2.1 gfaABC1D-cyto-GCaMP6f (Addgene viral prep # 52925-AAV5) was injected at 50 nL/min using a hydraulic injection apparatus driven by a syringe pump (UltraMicroPump, WPI, Sarasota, FL). Following the viral injection, a circular craniotomy (3 mm diameter) was centered at the stereotaxic coordinates (1.5 mm anterior and 1.8 mm lateral to bregma) using a trephine drill. The dura was left intact, and a custom chronic cranial window for mesoscopic two-photon imaging was placed above the craniotomy and secured using cyanoacrylate glue. A custom titanium head-plate was attached to the skull using cyanoacrylate glue. Finally, the components were secured using dental cement (C&B Superbond; Sun Medical). All the animals received an intraperitoneal bolus of antibiotic (BAYTRIL, Bayer, Germany) to prevent postsurgical infections.

### Mesoscale two-photon imaging in awake head-restrained mice

A two-photon random access mesoscope (2P-RAM, ^48^, ThorLabs Mesoscope, Thorlabs, Newton, NJ) was coupled with a pulsed laser source (Chameleon Ultra, 80 MHz repetition rate tuned at 920 nm, Coherent). Group delay dispersion was compensated using a prism-based compensation unit. 2P-RAM scanning unit was composed of a resonant scanner (24 kHz, CRS 12 K, Cambridge Technology), a pair of galvanometric mirrors, and an acoustic coil remote focusing unit. Image acquisition was controlled using Scanimage ^81^ (MBF Bioscience, Ashburn VA). Imaging was performed using a 0.6 NA objective (S/N:126, Jenoptik, Jena), and emitted fluorescence was collected using GaAsP photomultiplier tubes (PMT2103, Thorlabs, Newton, NJ) after band-pass filtering (520/70 nm). Laser beam intensity was adjusted using a Pockel cell (ConOptics Inc, Danbury). Laser beam power at the objective outlet was 50-70 mW. Imaging was conducted on a ∼1.5×1.5 mm field of view scanned using three tiled regions of interest of 1500×500 pixels (1 µm/pixel) resulting in a composite field of view of 1500×1500 pixels sampled at ∼3 Hz frame rate. T-series were motion corrected using an open-source implementation of up-sampled phase cross-correlation ^71, 80^ and the t-series median projection was used as reference frame. Three weeks after surgery the animals were handled by the operator for two days and habituated to the imaging setup. Starting from the second session, the animals were head-restrained for a progressively increasing amount of time, reaching 1 hour in approximately one week. Mice were free to run on a custom 3D printed wheel and water rewards were provided by the operator through a custom waterspout. Experimental sessions lasted between 1-1.5 hours. After each session, animals were returned to their home cages.

### Algorithm Open-source implementation and Datasets availability

ASTRA was developed in Python 3.6 ^82^ and PyTorch 1.2 ^83^ and the code is publicly available at (https://gitlab.iit.it/fellin-public/astra). ASTRA uses several open-source libraries like OpenCV ^63^, scikit-learn ^84^, scikit-image ^80^ and Scipy ^85^. The repository contains documentations, Docker (docker.com) image for fast installation, Jupyter notebook tutorials, bindings for widely used software (Fiji, ^72^ and MATLAB (MathWorks)), visualization and analysis tools, and a message/discussion board. DNN weights for all the datasets used in this study are reported in the repository. The four datasets, including individual and consensus annotations, will be shared upon publication.

### Statistics

In statistical testing of detection and segmentation performance of ASTRA, we used a two-sided Wilcoxon rank test. When performing multiple comparisons between ASTRA and human users of detection and segmentation performance, we used Holm-Bonferroni method for post-hoc correction.

### Time Analysis

We analyzed the computational performances of ASTRA in terms of processing time for the various steps in the Inference Pipeline. We used the following computing architecture, a Linux based workstation (Ubuntu 18.04.3 LTS distribution) equipped with 20 Intel(R) Core (TM) i9-9900X CPU clocked @ 3.50GHz, 130 GB of RAM, and 3 INVIDIA GeForce RTX 2080Ti GPUs.

## Author contributions

JB, SC, SP, and TF conceived the project. JB developed deep learning and preprocessing algorithms. SC built the experimental set up. JB and SC performed analyses. SC and SR performed experiments. JB, SC, and SR annotated data. JB, SC, SP, and TF wrote the paper. SP and TF supervised the project and secured funding.

## SUPPLEMENTARY FIGURES

**Figure S1.**
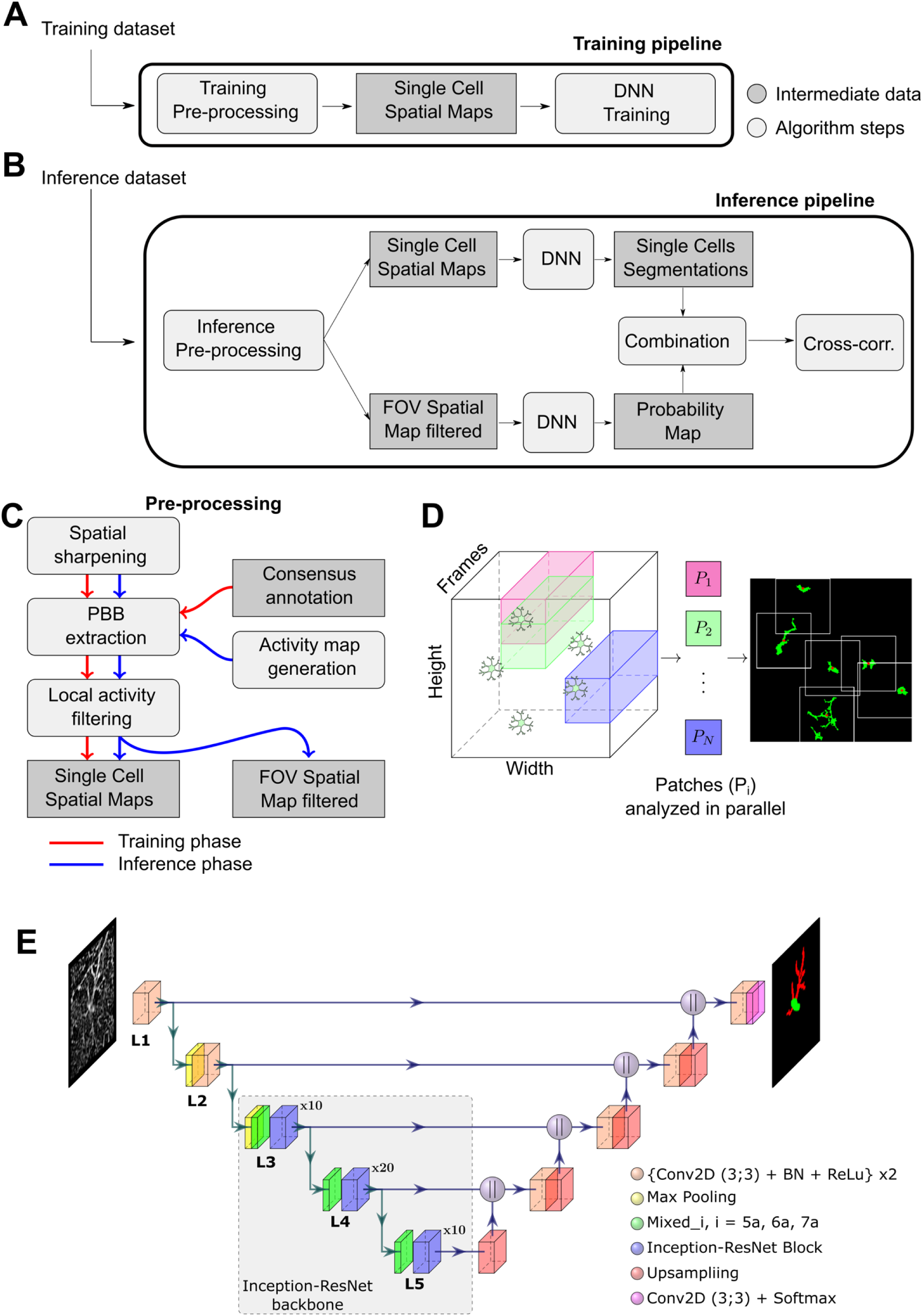
Schematic representation of ASTRA pipelines and modules. A-B) Flowcharts of the training (A) and inference (B) pipelines. C) Flowchart of the pre-processing steps: i) generation of the spatial sharpening map (Spatial Sharpening); ii) extraction of putative bounding boxes (PBB Extraction); and iii) local activity filtering (LA Filtering) of single-cell images. Please note that extraction of single-cell images during pre-processing of the training set relies on ground-truth segmentation. Extraction of single-cell images during pre-processing of the inference set relies on the activity map generator (Activity Map Generation). D) Schematic representation of ASTRA activity map generator: i) patch extraction; ii) patch parallel analysis; iii) clustering of active pixels. E) ASTRA DNN architecture. In each level Li with i = (1, 2, 3, 4, 5), height (H) and width (W) of the input image is reduced by a factor 2^i-1^. *Conv2D+BN+ReLu*: this block is composed of two consecutive sequences of 3 x 3 convolutional filters (Conv2D) followed by batchnorm normalization (BN) and rectified linear unit (Relu). *Max Pooling*: we used a kernel_size of (2,2) - the size of the sliding window where the maximum value of the input tensor is taken - resulting in input tensor of dimensions H and W reduced to H/2 and W/2. *Mixed_i*: in L3, we used Mixed_5a, in L4 we used Mixed_6a, and, in L5, we used Mixed_7a from Inception-ResNetv2 implementation in ^64^. *Inception-ResNet Block*: in L3 the block is composed as (Inception-ResNet-A, Block35)x10, in L4 the block is composed by (Inception-ResNet-B, Block 17)x20 and in L5 the block is composed by (Inception-ResNet-C, Block8)x10 from Inception-ResNetv2 implementation in (Szegedy et al. 2017). *Upsampling*: we adopted dense upsampling convolution (DUC, ^67^) to perform the upsampling on the input tensor. The input tensor dimensions are H x W x D and they are transformed to (2H) x (2W) x (D/4). *Conv2D+Softmax*: this block is composed by a 3×3 convolutional filter and a Softmax transformation.

**Figure S2.**
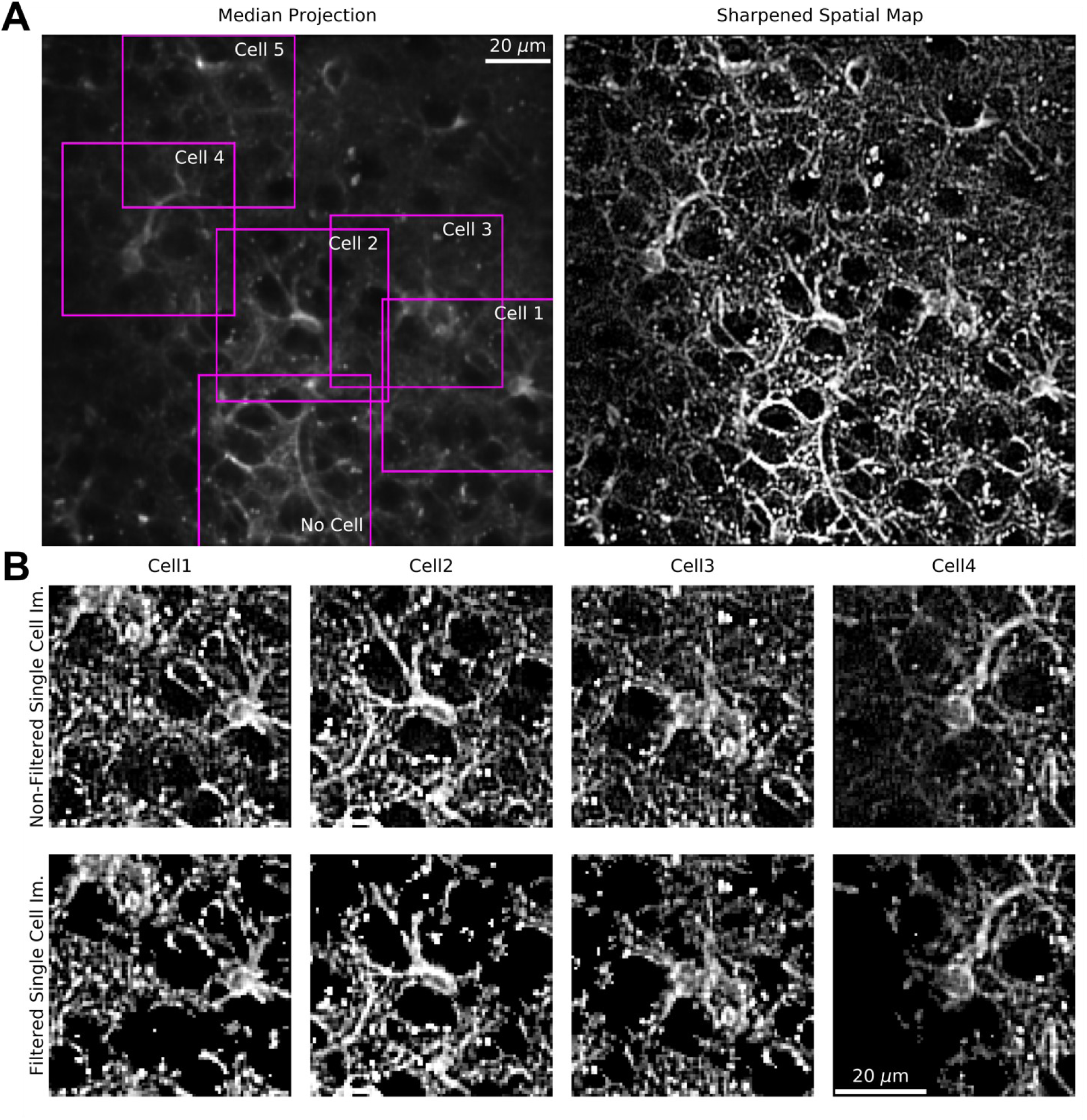
ASTRA pre-processing. A) Left: median projection of a representative FOV (Id:2) overlaid with putative bounding boxes computed by activity map generation. Right: spatial sharpening of the same FOV shown on the left panel. B) Top: zoom in showing sharpened images of four cells (cell 1-4) extracted from the putative bounding boxes shown in the left panel of A. Bottom: for each image the result of local activity filtering is shown.

**Figure S3.**
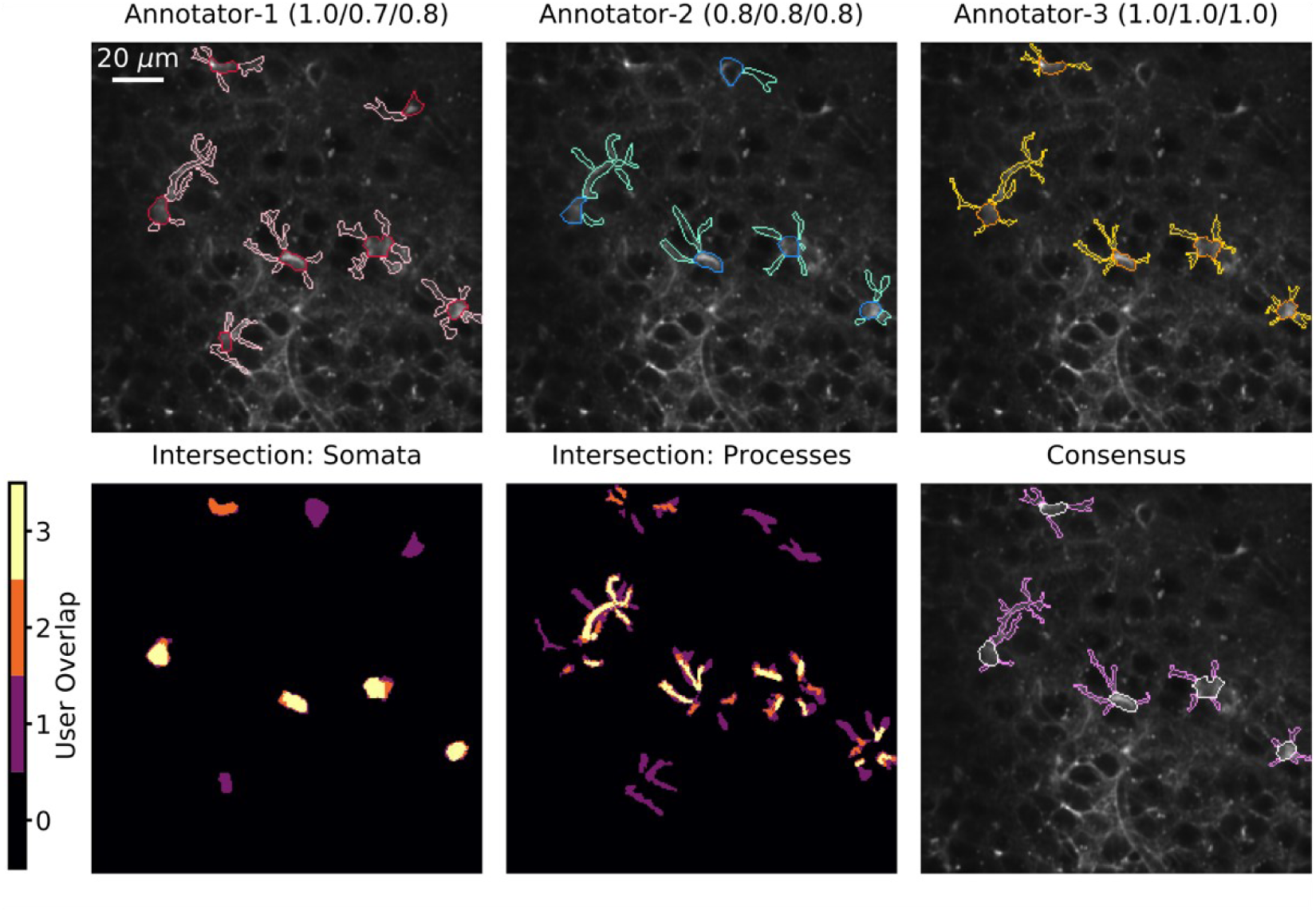
Generation of the consensus annotation. Top: individual manual annotations (colored contours) for FOV (Id:2) by three graders (annotator-1, annotator-2, and annotator-3). Manual annotations are plotted on top of the median projection of the two-photon t-series. The numbers in parenthesis in the top label report detection Precision, Recall, and F1 score. Bottom: intersection of somata annotations (left), intersection of process annotations (middle), and result of the consensus annotation (right).

**Figure S4.**
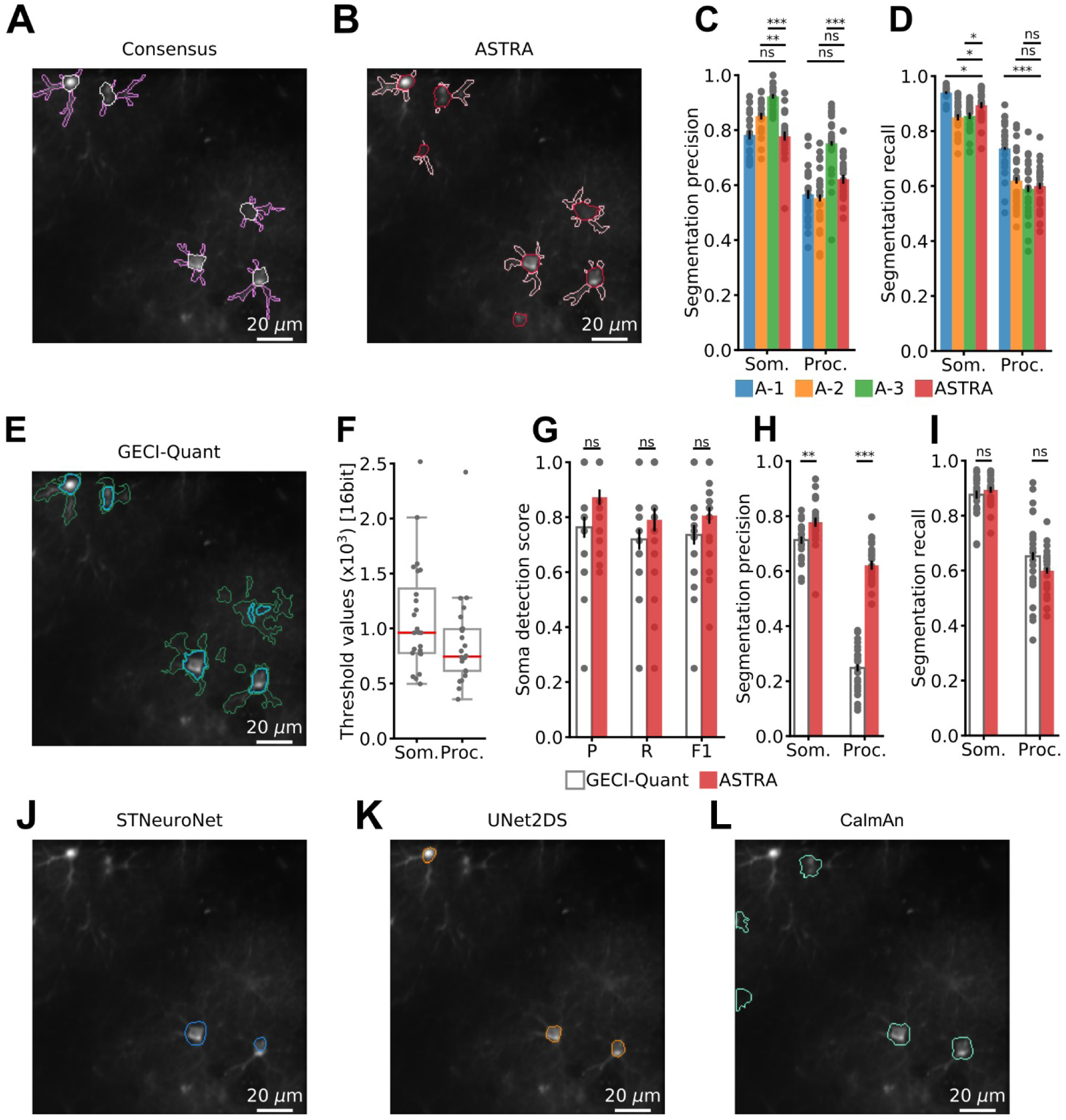
Benchmarking ASTRA against human annotators and against state-of-the-art cell detection and segmentation methods. A-B) Precision (A) and Recall (B) of somata and process segmentation for the three annotators and ASTRA (two-sided Wilcoxon rank sum test N = 24; LOOCV results). See also table S1. C-I) ASTRA semantic segmentation against GECI-quant segmentation. Representative example of segmentations of somata and processes for: C) the consensus annotation, somata (white), processes (light purple); D) ASTRA, somata (red), processes (pink); E) GECI-Quant, somata (light blue), processes (green). F) GECI-Quant user defined thresholds distributions for dataset-1. Box charts show the median values (red line) and the interquartile range (IQR, black top and bottom limit of the box). The whiskers extend to 1.5 times the IQR. G) GECI-Quant soma detection vs. ASTRA in dataset-1. Precision, Recall, and F1-score are shown (two-sided Wilcoxon signed rank sum test, N = 24; LOOCV results). H-I) Precision (H) and Recall (I) for somata and process segmentation (two-sided Wilcoxon rank sum test, N = 24; LOOCV results). See also table S2. J-L) Representative examples of **somata** segmentations on the same FOV shown in C for: J) STNeuroNet, somata (blue); K) UNet2DS somata (orange); L) CaImAn, somata (light green). In A-B and G-H-I: n.s., not significant, * p < 0.05, ** p < 0.005, and *** p < 0.0005.

**Figure S5.**
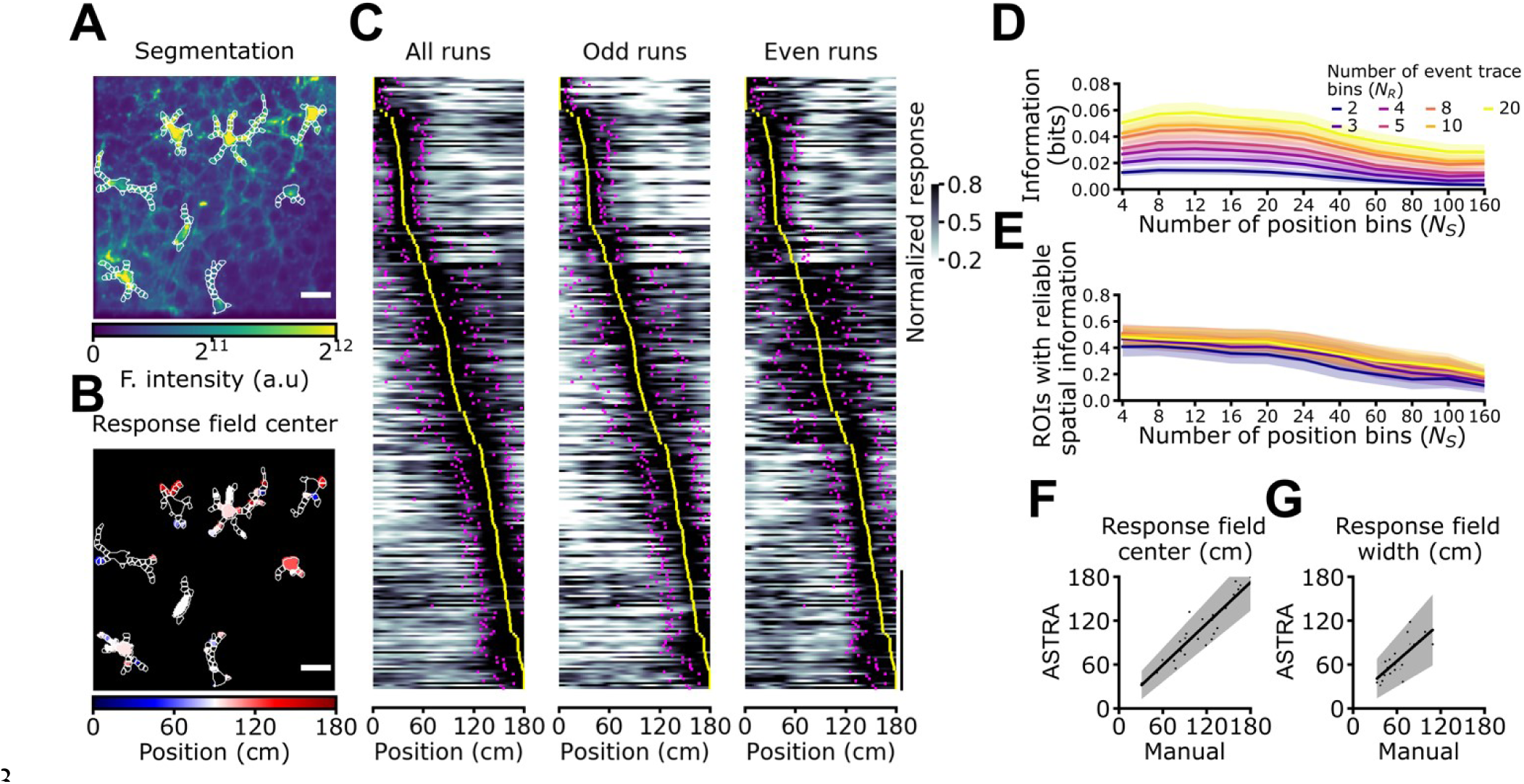
ASTRA automated analysis replicates previously published results obtained with manual annotation. A) ASTRA segmentation (white) of GCaMP6f-labeled astrocytes in the CA1 pyramidal layer. Experimental data from ^15^. B) Same FOV as in (A) with astrocytic ROIs color-coded according to response field position along the virtual corridor. In (A) and (B) scale bar: 20 μm. C) Normalized astrocytic calcium responses as a function of position for astrocytic ROIs that contain significant spatial information (N = 260 ROIs with reliable spatial information out of 595 total ROIs, 7 imaging sessions from 3 animals, see Methods for details). Responses are ordered according to the position of the center of the response field (from minimum to maximum). Left panel, astrocytic calcium responses from all trials. Center and right panels, astrocytic calcium responses from odd (center) and even (right) trials. Yellow dots indicate the center position of the response field, while magenta dots indicate the extension of the field response (see Methods, vertical scale: 50 ROIs). D-E) Bias-corrected mutual information values (D) and fraction of ROIs encoding reliable spatial information (E) as a function of the number of bins for the stimulus (animal position along the linear track). Colors indicate binning of the response (calcium event trace). F-G) Cell-wise comparison of average response field center position (F) and width (G) between the results obtained with ASTRA (y axis) and those reported in ^15^ (x axis). Dots represent average of ROI parameters for each cell. The black line is least-squares linear fit (in (F) y = 0.94x+2.79 cm, R^2^ = 0.87; in (G) y = 0.87x+12.41 cm, R^2^ = 0.56). In (F-G) n = 33 true positive astrocytes detected by ASTRA in 7 imaging sessions from 3 mice.

**Figure S6.**
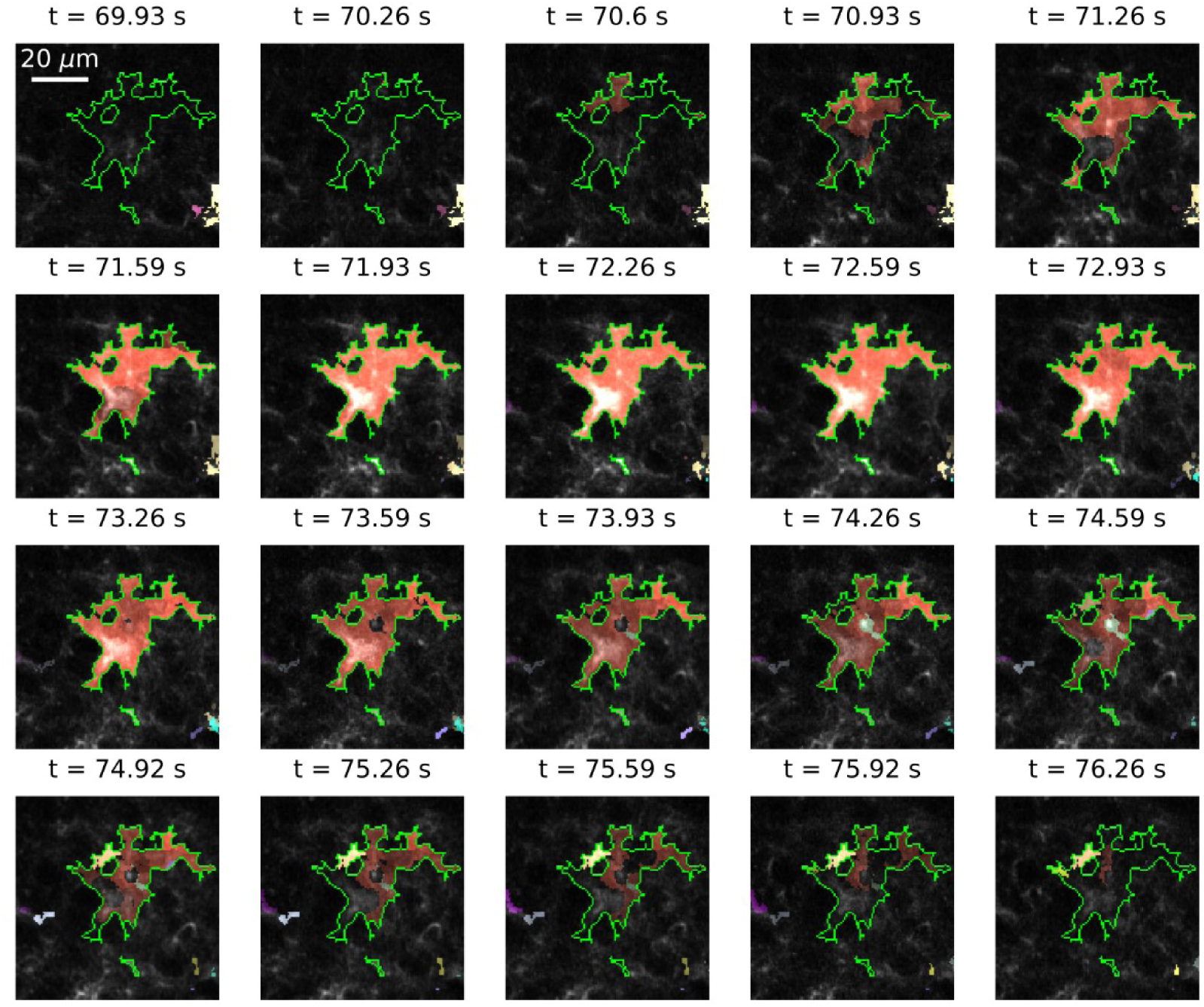
ASTRA seeding of event-based segmentation. Example of a spatiotemporal Ca^2+^ events (red to white colors) detected with AQuA when seeded with the astrocytic domain (green line) identified by ASTRA. Each image represents a single frame of a representative t-series (id: 2, dataset-1). Colors superimposed to each frame represent a detected event in the astrocyte. Frame acquisition time is reported on the top of each image.

**Figure S7.**
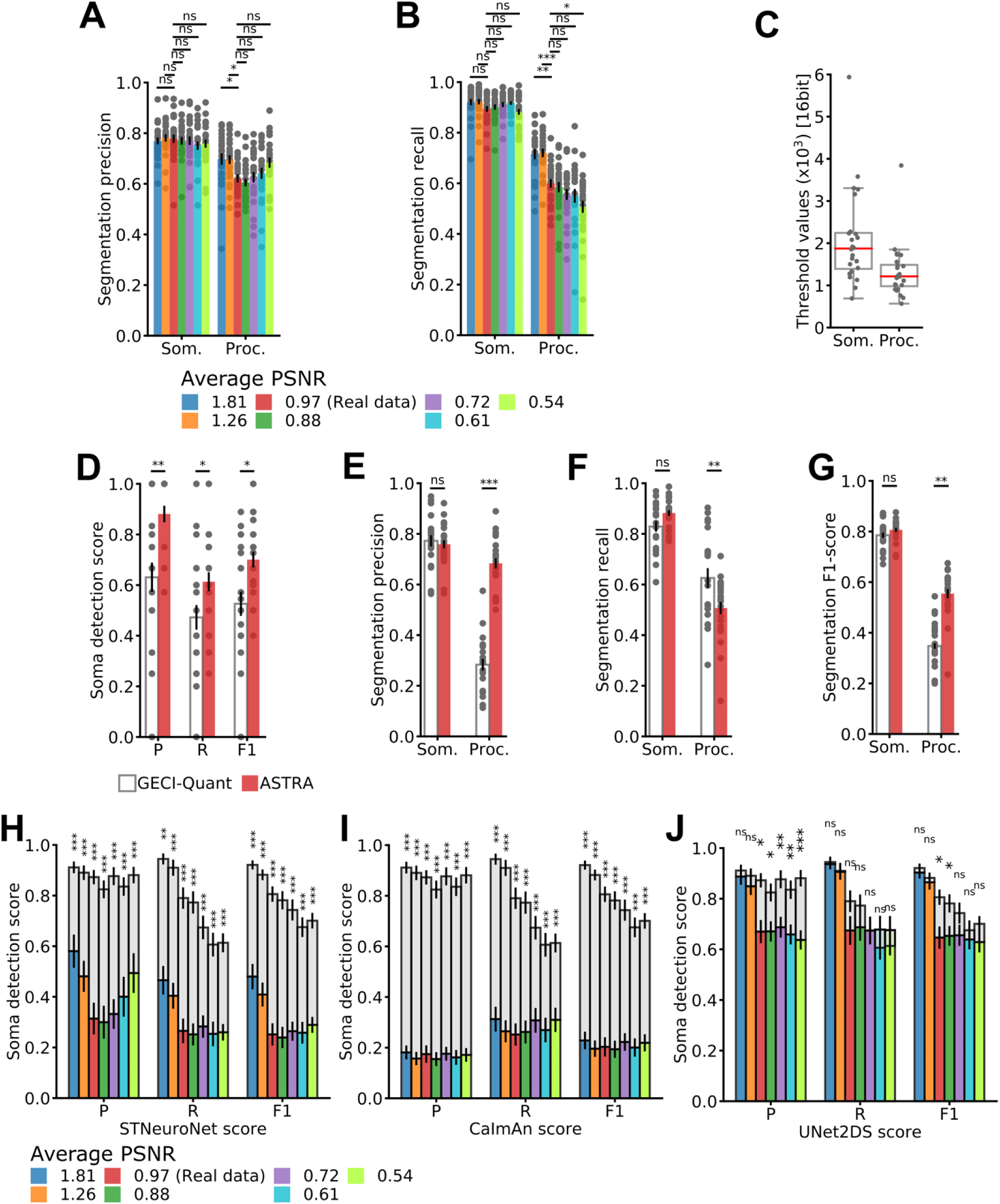
Comparative analysis of the effect of signal-to-noise ratio regimens on detection and segmentation performances. A-B) Precision (A) and Recall (B) for ASTRA segmentation of somata and processes under the different simulated conditions of PSNR (two-sided Wilcoxon rank sum test, N = 24; LOOCV results). C) Distribution of GECI-Quant thresholds for the 0.54 PSNR dataset for somata (Som.) and processes (Proc.). Box charts show the median values (red line) and the interquartile range (IQR, black top and bottom limit of the box). The whiskers extend to 1.5 times the IQR. D) Precision, Recall, and F1-score for soma detection for GECI-Quant (white) and ASTRA (red) for the 0.54 PSNR dataset (two-sided Wilcoxon signed rank sum test, N = 24; LOOCV results). E-G) Segmentation Precision (E), Recall (F), and F1-score (G) GECI-Quant (white) and ASTRA (red) for the 0.54 PSNR dataset (two-sided Wilcoxon rank sum test, N = 24; LOOCV results). H-J) Effect of artificial noise on soma detection performances. Detection Precision, Recall and F1-score for ASTRA (grey bars), STNeuronet (H), CaIman (I), and UNet2DS (J) on the same dataset under different regimens of PSNR (two-sided Wilcoxon rank sum test, N = 24; LOOCV results). In all panels: n.s., not significant, * p < 0.05, ** p < 0.005, and *** p < 0.0005.

**Figure S8.**
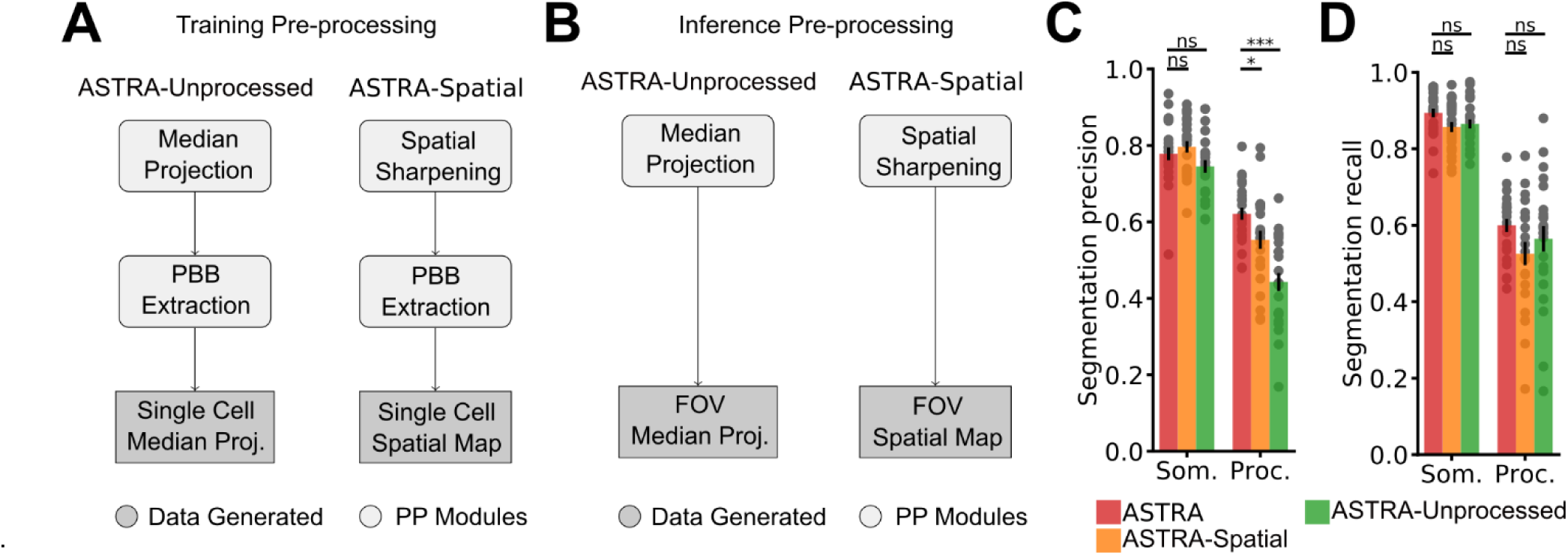
Impact of pre-processing on ASTRA performance. A) Flow-chart describing the pre-processing block in the training pipeline for ASTRA-Naïve and ASTRA-Spatial (see main results). In ASTRA-unprocessed, the DNN is trained with the single cell images extracted from the median projection of the FOVs. In ASTRA-Spatial, the DNN is trained with the single cell images extracted from the spatial map of the FOVs. B) Flow-chart of pre-processing block in the inference pipeline for ASTRA-unprocessed and ASTRA-Spatial. In ASTRA-unprocessed, the DNN directly evaluates median projection of the whole FOV. In ASTRA-Spatial, the DNN evaluates the spatial map of the whole FOV. C-D) Segmentation Precision (C) and Recall (D) for ASTRA-unprocessed, ASTRA-Spatial, and ASTRA on dataset-1 (two-sided Wilcoxon rank sum test, N = 24; LOOCV results). In C-D: n.s., not significant, * p < 0.05, ** p < 0.005, and *** p < 0.0005.

**Figure S9.**
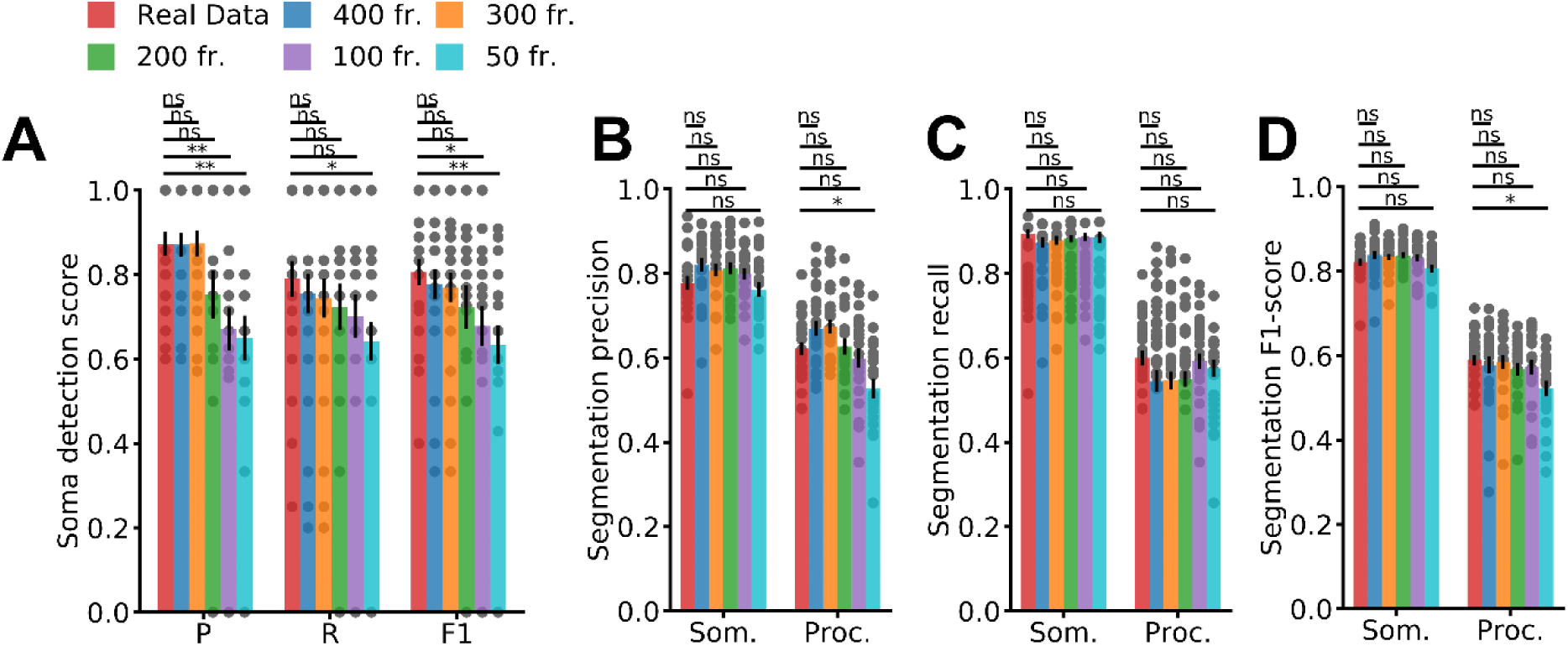
ASTRA performance as a function of recording length. A) ASTRA detection Precision, Recall, and F1-score for t-series of different length (two-sided Wilcoxon rank sum test, N = 24; LOOCV results). B-D) ASTRA segmentation Precision (B), Recall (C), and F1-score (D) for t-series of different length (two-sided Wilcoxon rank sum test, N = 24; LOOCV results).

**Figure S10.**
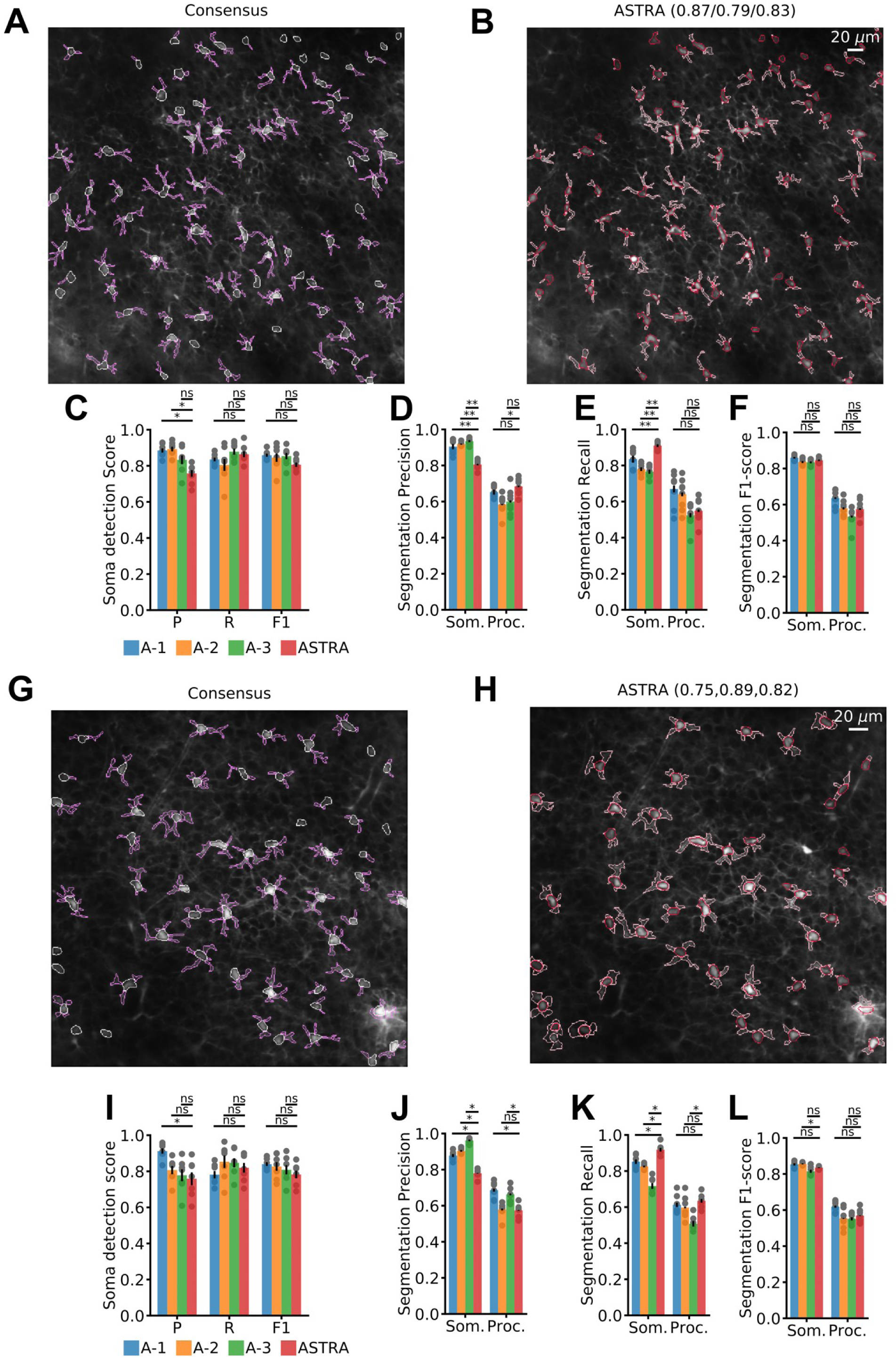
ASTRA detection and segmentation performance on dataset 2 and 3. A) Consensus annotation of one representative FOV (Id: 3) showing Td-Tomato-expressing astrocytes. B) ASTRA segmentation result for the same FOV shown in (A). C) Detection Precision, Recall, and F1-score for ASTRA and the three annotators (two-sided Wilcoxon rank sum test, N = 8; LOOCV results). D-F) Segmentation Precision, Recall, and F1-score of ASTRA and the three annotators for somata (Som.) and processes (Proc.) (two-sided Wilcoxon rank sum test, N = 8; LOOCV results). See also Table S3. G) Consensus annotation of one representative FOV (Id: 5) showing Td-Tomato-expressing astrocytes. H) ASTRA segmentation result for the same FOV shown in (G). I) Detection Precision, Recall, and F1-score for ASTRA and the three annotators (two-sided Wilcoxon rank sum test N = 7; LOOCV results). J-L) Segmentation Precision, Recall, and F1-score of ASTRA and the three annotators for somata (Som.) and processes (Proc.) (two-sided Wilcoxon rank sum test, N=7; LOOCV results). In C-F and I-L: n.s., not significant, * p < 0.05, ** p < 0.005, and *** p < 0.0005. See also Table S4 and Table S5.

**Figure S11.**
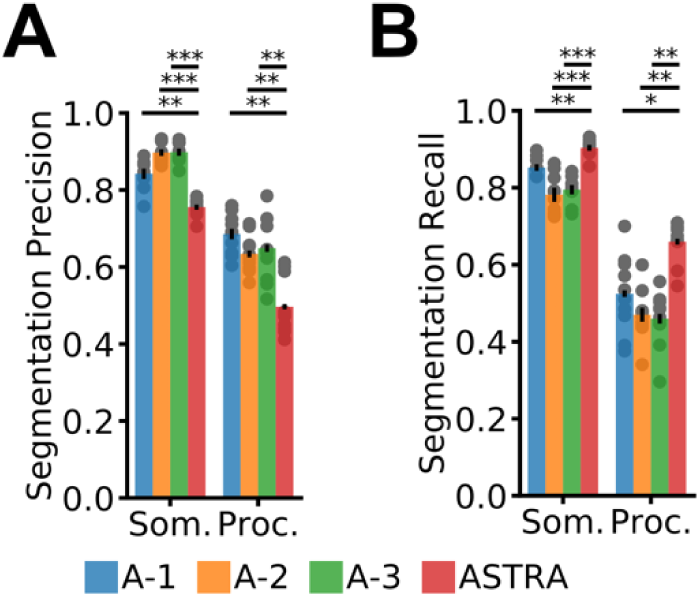
ASTRA detection and segmentation performance on dataset-4. A-B) Segmentation Precision (A) and Recall (B) of ASTRA and the three annotators for somata (Som.) and processes (Proc.) (two-sided Wilcoxon rank sum test, N=10; LOOCV results). In all panels: n.s., not significant, * p < 0.05, ** p < 0.005, and *** p < 0.0005. See also Table S6.

**Figure S12.**
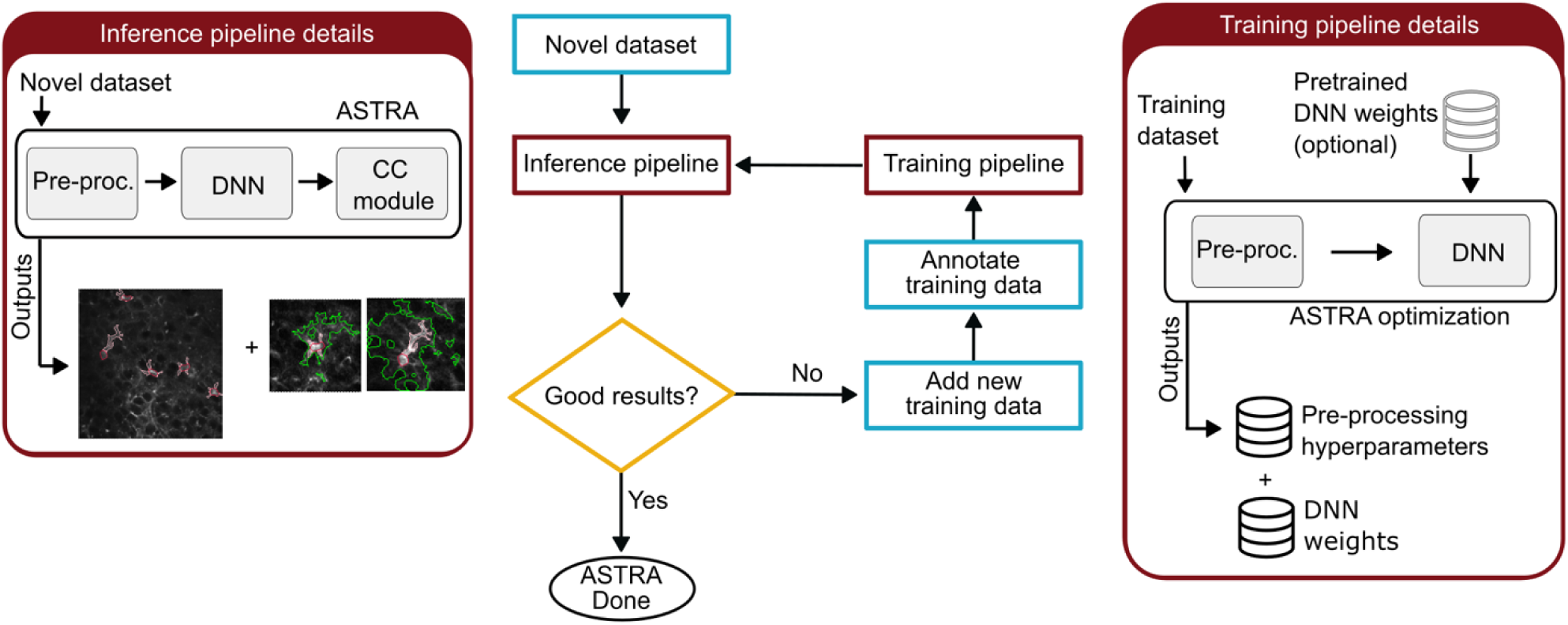
ASTRA workflow. The diagram delineates the workflow which is color-coded to identify input of new data and eventually new annotated data (cyan), ASTRA training and inference pipelines (dark red), and user evaluation of ASTRA results (yellow). Inference and training pipelines details are reported in the 2 boxes placed on the sides.

**Figure S13.**
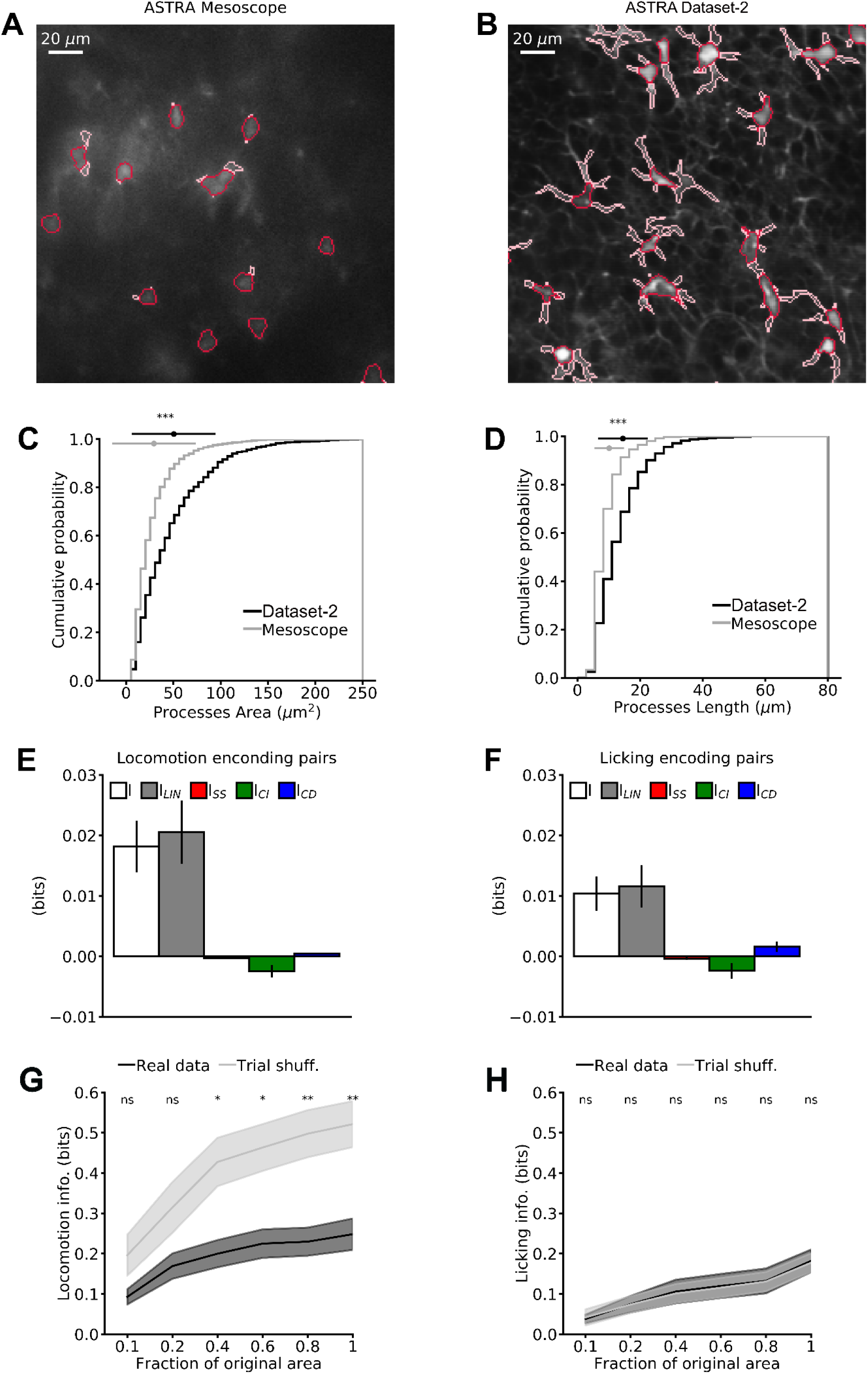
**Stimulus encoding properties of mesoscale astrocytic networks in the mouse neocortex**. A) Representative ASTRA somata and processes segmentation on a zoomed in image of 0.48 mm^2^ area extracted from a 2.25 mm^2^ area FOV recorded with the two-photon mesoscope. B) Representative ASTRA somata and processes segmentation on a FOV recorded with conventional two-photon configuration (Id: 3, in dataset-2). The FOV in B has the same size as in A. C-D) Cumulative distribution of area (C) and maximum length (D) of process ROIs segmented using ASTRA on the mesoscope dataset and on dataset-2. ROI area: mean ± std 29 ± 23 µm^2^ for mesoscope dataset and 51 ± 44 µm^2^ for dataset-2, t-test N = 964 ROIs from 15 imaging sessions and N = 1483 ROIs from 8 imaging sessions for the mesoscope dataset and dataset-2, respectively. Maximal ROI length: mean ± std 10 ± 5 um for mesoscope dataset and 15 ± 8 um for dataset-2, t-test N = 964 ROIs from 15 imaging sessions and N = 1483 ROIs from 8 imaging sessions for the mesoscope dataset and dataset-2, respectively. E-F) Information breakdown analysis from ROI pairs. E) Mutual information about locomotion (I, white) encoded by a pair of ROIs, sum of the mutual information encoded in the response of each member of the pair (ILIN, grey), the information component due to signal-similarity (ISS, red), stimulus independent information contribution of correlation (ICI, green), and stimulus dependent information contribution of correlation (ICD, blue) are shown. Data are represented as mean ± sem and were collected in 13 imaging sessions from 3 animals for running encoding pairs. F) Same as in (E) but for encoding of licking behavior. Data from 8 imaging sessions from 3 animals. G-H) Decoded information increases with the spatial size of the network. G) Decoded information of locomotion from astrocytic population vectors on real (black), and trial-shuffled (gray) as a function of the fraction of the original FOV area. For each fraction of area two-sided Wilcoxon signed rank test (N = 13) has been performed between real and trial-shuffled data. H) Decoded information of licking from astrocytic population vectors on real (black), and trial-shuffled (gray) as a function of the fraction of the original FOV area. For each fraction of area two-sided Wilcoxon signed rank test (N = 8) has been performed between real and trial-shuffled data. Data are represented as mean ± sem. In panel A data are obtained from 13 imaging sessions in 3 animals. In panel B data are obtained from 8 imaging sessions in 3 animals.

**Figure S14.**
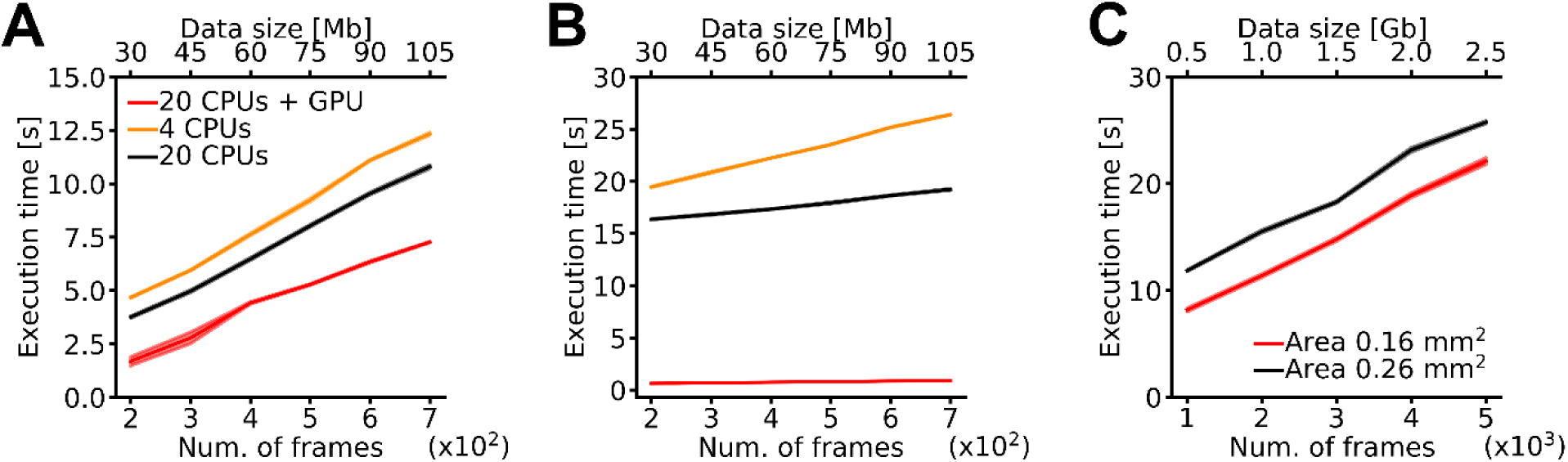
Execution time of the ASTRA inference pipeline. A) Execution time of ASTRA inference pipeline without the cross-correlation analysis as a function of the size of the input t-series. The different colors indicate the execution time for three different hardware configurations: 4 CPUs, 20 CPUs, and 20 CPUs + GPU. B) Execution time for the astrocytic domain module as a function of the size of the input t-series. Color code same as in (A). Please note that the GPU configuration is faster than the multi-processing CPU configuration. This is because the computation of cross-correlation value between pixels can be massively parallelizable with GPUs. C) Execution time for the inference pipeline without cross-correlation analysis as a function of the size of the input t-series for dataset-2 (black, area 0.26 mm^2^) and dataset-3 (red, area of FOV 0.16 mm^2^).

**Table S1.** Results of each annotator against consensus annotation of dataset-1. Detection and Segmentation results are given using F1-score (Precision, Recall) metrics (mean ± sem).

|  | Detection | Segmentation |  |
| --- | --- | --- | --- |
|  |  | Somata | Processes |
| **Annotator-1** | 0.88±0.02<br>(0.91±0.02,0.89±0.03) | 0.845±0.009<br>(0.78±0.02,0.939±0.005) | 0.62±0.02<br>(0.56±0.02,0.74±0.01) |
| **Annotator -2** | 0.88±0.02<br>(0.87±0.03,0.91±0.02) | 0.84±0.01<br>(0.85±0.01,0.852±0.006) | 0.56±0.01<br>(0.55±0.02,0.62±0.02) |
| **Annotator -3** | 0.90±0.02<br>(0.89±0.03,0.92±0.02) | 0.882±0.007<br>(0.923±0.008,0.86±0.01) | 0.64±0.02<br>(0.75±0.02,0.59±0.02) |

**Table S2.** Results of ASTRA, STNeuronet, Caiman and GECI-Quant against consensus annotation of dataset-1. Detection and Segmentation results are given using F1-score (Precision, Recall) metrics (mean ± sem)

|  | Detection | Segmentation |  |
| --- | --- | --- | --- |
|  |  | Somata | Processes |
| **ASTRA** | 0.81±0.04<br>(0.79±0.04,0.87±0.03) | 0.822±0.008<br>(0.78±0.02,0.89±0.01) | 0.59±0.01<br>(0.62±0.02,0.60±0.01) |
| **STNeuronet** | 0.27±0.05<br>(0.26±0.05,0.32±0.06) | - | - |
| **Caiman** | 0.20±0.04<br>(0.25±0.04,0.17±0.03) | - | - |
| **UNet2DS** | 0.65±0.04<br>(0.67±0.06,0.67±0.05) | - | - |
| **GECI-Quant** | 0.74±0.04<br>(0.72±0.04,0.76±0.04) | 0.775±0.008<br>(0.72±0.01,0.88±0.01) | 0.33±0.02<br>(0.25±0.02,0.65±0.03) |

**Table S3.** Results of ASTRA, and AQuA in reconstructing astrocytic morphology. Results are F1-score (Precision, Recall) metrics (mean ± sem) vs consensus annotation.

|  | **F1-score**<br>**(Precision, Recall)** |
| --- | --- |
| **ASTRA** | 0.62±0.03<br>(0.61±0.03,0.65±0.03) |
| **AQuA** | 0.23±0.02<br>(0.12±0.02,0.53±0.2) |

**Table S4.**
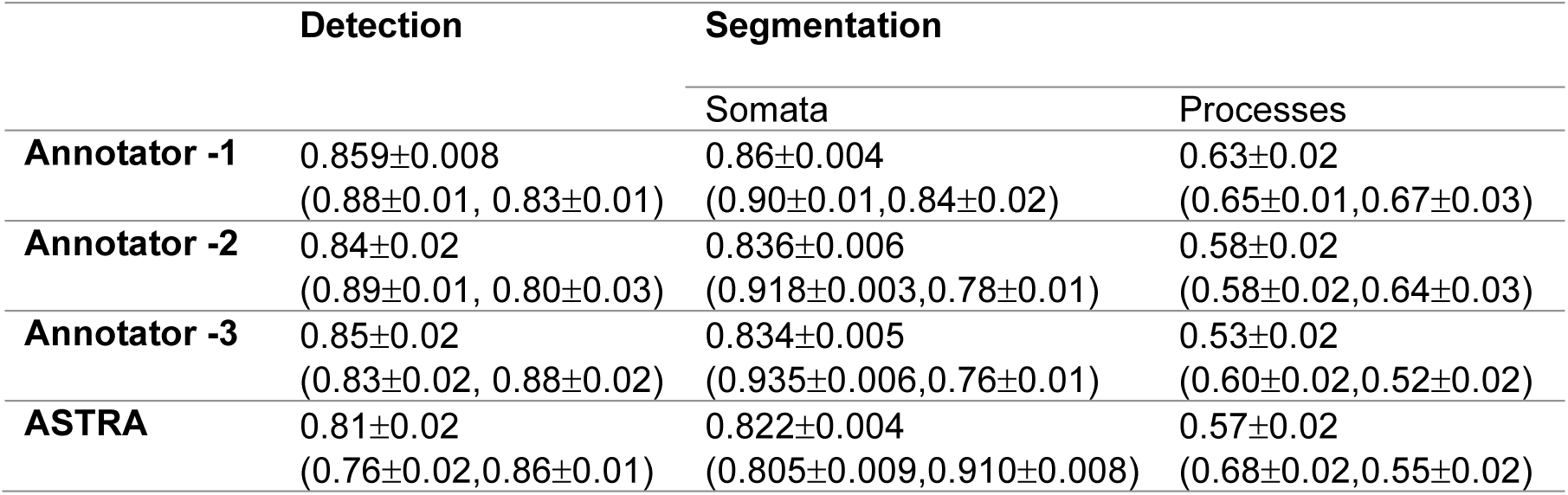
Results of each annotator and ASTRA against consensus annotation of dataset-2. Detection and Segmentation results are given using F1-score (Precision, Recall) metrics (mean ± sem)

|  | Detection | Segmentation |  |
| --- | --- | --- | --- |
|  |  | Somata | Processes |
| **Annotator -1** | 0.859±0.008<br>(0.88±0.01, 0.83±0.01) | 0.86±0.004<br>(0.90±0.01,0.84±0.02) | 0.63±0.02<br>(0.65±0.01,0.67±0.03) |
| **Annotator -2** | 0.84±0.02<br>(0.89±0.01, 0.80±0.03) | 0.836±0.006<br>(0.918±0.003,0.78±0.01) | 0.58±0.02<br>(0.58±0.02,0.64±0.03) |
| **Annotator -3** | 0.85±0.02<br>(0.83±0.02, 0.88±0.02) | 0.834±0.005<br>(0.935±0.006,0.76±0.01) | 0.53±0.02<br>(0.60±0.02,0.52±0.02) |
| **ASTRA** | 0.81±0.02<br>(0.76±0.02,0.86±0.01) | 0.822±0.004<br>(0.805±0.009,0.910±0.008) | 0.57±0.02<br>(0.68±0.02,0.55±0.02) |

**Table S5.** Results of each annotator and ASTRA against consensus annotation of dataset-3. Detection and Segmentation results are given using F1-score (Precision, Recall) metrics (mean ± sem)

|  | Detection | Segmentation |  |
| --- | --- | --- | --- |
|  |  | Somata | Processes |
| **Annotator -1** | 0.84±0.01<br>(0.91±0.01, 0.78±0.02) | 0.853±0.006<br>(0.878±0.008,0.85±0.01) | 0.62±0.01<br>(0.69±0.02,0.61±0.02) |
| **Annotator -2** | 0.83±0.02<br>(0.81±0.02, 0.85±0.03) | 0.856±0.003<br>(0.904±0.005,0.825±0.005) | 0.55±0.02<br>(0.58±0.01,0.60±0.02) |
| **Annotator -3** | 0.81±0.03<br>(0.78±0.03, 0.85±0.03) | 0.815±0.009<br>(0.962±0.005,0.717±0.02) | 0.55±0.01<br>(0.66±0.01,0.51±0.01) |
| **ASTRA** | 0.78±0.02<br>(0.76±0.04,0.82±0.03) | 0.835±0.002<br>(0.780±0.008,0.92±0.01) | 0.57±0.01<br>(0.57±0.02,0.63±0.01) |

**Table S6.** Results of each annotator and ASTRA against consensus annotation of dataset-4. Detection and Segmentation results are given using F1-score (Precision, Recall) metrics (mean ± sem)

|  | Detection | Segmentation |  |
| --- | --- | --- | --- |
|  |  | Somata | Processes |
| **Annotator -1** | 0.81±0.01<br>(0.90±0.01, 0.75±0.02) | 0.835±0.006<br>(0.84±0.01,0.852±0.007) | 0.55±0.02<br>(0.69±0.02,0.52±0.03) |
| **Annotator -2** | 0.72±0.02<br>(0.73±0.03, 0.73±0.03) | 0.827±0.007<br>(0.897±0.007,0.78±0.01) | 0.50±0.02<br>(0.63±0.01,0.47±0.02) |
| **Annotator -3** | 0.74±0.03<br>(0.70±0.06, 0.80±0.01) | 0.834±0.004<br>(0.898±0.008,0.79±0.01) | 0.50±0.02<br>(0.65±0.03,0.46±0.02) |
| **ASTRA** | 0.80±0.02<br>(0.78±0.03,0.82±0.02) | 0.813±0.006<br>(0.755±0.007,0.904±0.006) | 0.53±0.01<br>(0.50±0.02,0.66±0.02) |

**Table S7.** ASTRA DNN training parameters

|  |  | Epochs |  | Optimizer | lr | Batch size | Input image size |
| --- | --- | --- | --- | --- | --- | --- | --- |
|  |  | N1 | N2 |  |  |  |  |
| **Dataset-1** | | 12 | 3 | Adam | $10^{-4}$ | 35 | 96x96 |
| **Dataset-2** | | 12 | 3 | Adam | $10^{-4}$ | 35 | 48x48 |
| **Dataset-3** | Training. on Dataset-1 | - | - | - | - | - | - |
| **Dataset-4** | Training on Dataset-1 | - | - | - | - | - | - |

